# Direct-fit to nature: an evolutionary perspective on biological (and artificial) neural networks

**DOI:** 10.1101/764258

**Authors:** Uri Hasson, Samuel A. Nastase, Ariel Goldstein

**Affiliations:** Princeton Neuroscience Institute, Princeton University, Princeton, NJ, USA; Department of Psychology, Princeton University, Princeton, NJ, USA

**Keywords:** evolution, experimental design, interpolation, learning, neural networks

## Abstract

Evolution is a blind fitting process by which organisms, over generations, adapt to the niches of an ever-changing environment. Does the mammalian brain use similar brute-force fitting processes to learn how to perceive and act upon the world? Recent advances in training deep neural networks has exposed the power of optimizing millions of synaptic weights to map millions of observations along ecologically relevant objective functions. This class of models has dramatically outstripped simpler, more intuitive models, operating robustly in real-life contexts spanning perception, language, and action coordination. These models do not learn an explicit, human-interpretable representation of the underlying structure of the data; rather, they use local computations to interpolate over task-relevant manifolds in a high-dimensional parameter space. Furthermore, counterintuitively, over-parameterized models, similarly to evolutionary processes, can be simple and parsimonious as they provide a versatile, robust solution for learning a diverse set of functions. In contrast to traditional scientific models, where the ultimate goal is interpretability, over-parameterized models eschew interpretability in favor of solving real-life problems or tasks. We contend that over-parameterized blind fitting presents a radical challenge to many of the underlying assumptions and practices in computational neuroscience and cognitive psychology. At the same time, this shift in perspective informs longstanding debates and establishes unexpected links with evolution, ecological psychology, and artificial life.

## Introduction

On a moment-to-moment basis, the brain is assimilating dynamic, multidimensional information about the world in order to produce rich, context-dependent behaviors. Confronted with such complexity, experimental neuroscientists traditionally design controlled experiments to reduce the dimensionality of the problem to a few factors conceived by the experimenter (Fisher, 1935). This reductionist program relies on a core commitment to the assumption that the neural computations supporting many of our cognitive functions can be decontextualized and decomposed into a handful of latent features; that these features are human-interpretable and can be manipulated in isolation; and that the piecemeal recomposition of these features will yield a satisfying understanding of brain and behavior.

In parallel to the research in neuroscience and psychology laboratories, artificial neural network (ANN) models are attaining human-level behavioral performance across many tasks, such as face recognition (e.g., Taigman et al., 2014), language processing (e.g., Radford et al., 2019), complex gameplay (e.g., Jaderberg et al., 2019), and motor learning (e.g., Levine et al., 2018). This research program effectively abandoned traditional experimental design and simple interpretable models, instead putting a premium on behavior (i.e., task performance) and embracing complexity. Such models learn how to recognize faces or respond to natural-language inquiries directly from the structure of the real world by optimizing millions of parameters (“big” models) over millions of examples (“big” data; LeCun et al., 2013). While the use of ANNs to model cognitive processes can be traced back through connectionism and parallel distributed processing (PDP), modern neural networks also substantially diverge from the tendency of classical connectionist modeling to rely on relatively small-scale, interpretable models with well-controlled inputs (e.g., Rumelhart et al., 1986; McClelland and Rogers, 2003).

In this paper, we consider how ANNs learn to perform complicated cognitive tasks, and whether the solution is at all relevant to cognitive neuroscientists. We use face recognition and language processing as examples of cognitive tasks, which have been extensively studied in cognitive neuroscience (see Box 2). Hundreds of experimental manipulations have been used to probe the neural machinery supporting face recognition and language processing, each aiming to isolate a handful of relevant factors underlying such functions. While as a field we have had great success in identifying neural variables that covary with our experimental variables, we are still far from understanding the neural computations that support such behaviors in real-life contexts and our toy models generally cannot compete with ANNs. Cognitive neuroscientists traditionally advocate for a privileged role of behavior in constraining models of neural information processing (Krakauer et al., 2017). We agree, with the caveat that contrived experimental manipulations may not provide sufficiently rich behavioral contexts for testing our models. We contend that advances in ANNs are the result of a strict adherence to the primacy of behavior and task performance, with the ambition (and commercial incentive) of building models that generalize to real-world contexts.

Similar to biological neural networks (BNNs), ANNs are trained to perform meaningful actions on real multidimensional data in real-life contexts. Across species and models, BNNs and ANNs can differ considerably in their circuit architecture, learning rules, and objective functions (Richards et al., 2019). All networks, however, use an iterative optimization process to pursue an objective given their input or environment—a process we refer to as “direct fit” (inspired by Gibson’s use of the term “direct perception”, as discussed below; Gibson, 1979). We draw on an analogy to the blind fitting processes observed in evolution by natural selection, and argue that ANNs and BNNs belong to the same family of direct-fit models. Both produce solutions that are mistakenly interpreted in terms of elegant design principles, but in fact reflect the interdigitation of “mindless” optimization processes and the structure of the world. This framework undercuts the assumptions of traditional experimental approaches, and makes unexpected contact with longstanding debates in developmental and ecological psychology. While direct-fit optimization provides the necessary foundations for many behaviors, current models still fall short on some high-level cognitive tasks. In the last section, we discuss limitations of this framework and the implications for high-level cognition.

### Simple versus multidimensional models

As with any scientific model, neuroscientific models are often judged based on their interpretability (i.e., providing an explicit, formulaic description of the underlying causes) and generalization (i.e., the capacity for prediction over broad, novel contexts; e.g., von Neumann, 1955^1^). However, in practice, interpretability and generalization are often at odds: interpretable models may have considerable explanatory appeal but poor predictive power, while high-performing predictive models may be difficult to interpret (Breiman, 2001; Shmueli, 2010; Yarkoni and Westfall, 2017).

This tension is particularly striking when modeling brain and behavior. The brain itself, in orchestrating behavior, is by conventional standards a wildly over-parameterized modeling organ (Conant and Ashby, 1970). Each cubic millimeter of cerebral cortex contains roughly 50,000 neurons that may support on the scale of 6,000 adjustable synapses with their neighboring and distant cells. This yields a staggering number of about 300 million adjustable parameters in each cubic millimeter of cortex, and over 100 trillion adjustable synapses across the entire brain (Azevedo et al., 2009; Kandel et al, 2013). This over-parameterized modeling organ is an evolutionary solution for producing flexible, adaptive behavior in a complex world.

In contrast, neuroscientists often reduce the complexity of the task (or stimulus) by using low-dimensional experimental manipulations in hopes of increasing the interpretability of observed neural processes. By analyzing the neural responses in such controlled situations, neuroscientists search the brain for simple latent factors for describing the code that underlies a neural computation. These experimental manipulations are often inspired by our “folk” or phenomenological understanding of the mind, brain, or world, and in turn, yield results reflecting our own assumptions (Meehl, 1990; Rozenblit and Keil, 2002; Jolly and Chang, 2018). That is, our simple models of the brain often boil down to models of our experimental design.

We are entering a new era in psychology and neuroscience in which over-parameterized models trained on big data are increasingly more powerful and dramatically outstrip simple, interpretable models in producing human-level “behavioral” performance across multiple cognitive tasks. While the power of over-parameterized models in machine learning is becoming apparent, there is fierce debate about whether they provide any insight as to the underlying neural code of biological organisms (e.g., Lake et al., 2017; Marcus, 2018a).

In this paper, we argue that neural computation is grounded in brute-force direct fitting, which relies on over-parameterized optimization algorithms to increase predictive power (generalization) without explicitly modeling the underlying generative structure of the world. We first differentiate two forms of generalization, extrapolation and interpolation^2^. Traditionally, interpolation was viewed as a weak form of generalization, due to its local (non-generative) nature. Here we argue that in the context of direct fit and big, real-world data, interpolation can provide a mindless, yet powerful, form of generalization (potentially eschewing the need for extrapolation).

### Interpolation and extrapolation

Statistics textbooks usually associate over-parameterized models with overfitting and contrast them with ideal-fit (also denoted as “appropriate fit” and “just-right fit”) and underfit models (Figure 1A-C). An underfit model is a model with too few parameters to capture the underlying structure of the observed data and thus provides poor prediction or generalization (Figure 1A). An overfit model is flexible enough to fit and/or memorize the structure of a training sample (including structureless noise and idiosyncrasies specific to the training set) to the extent that it fails to learn the structure needed for generalization (Figure 1C). An ideal-fit model is a model that learns the underlying generative or global structure of the data by exposing a few latent factors or rules (Figure 1B). As opposed to the underfit and the overfit models, the ideal-fit model is capable of generalization: accurately predicting new observations never seen during training.

**Figure 1.**
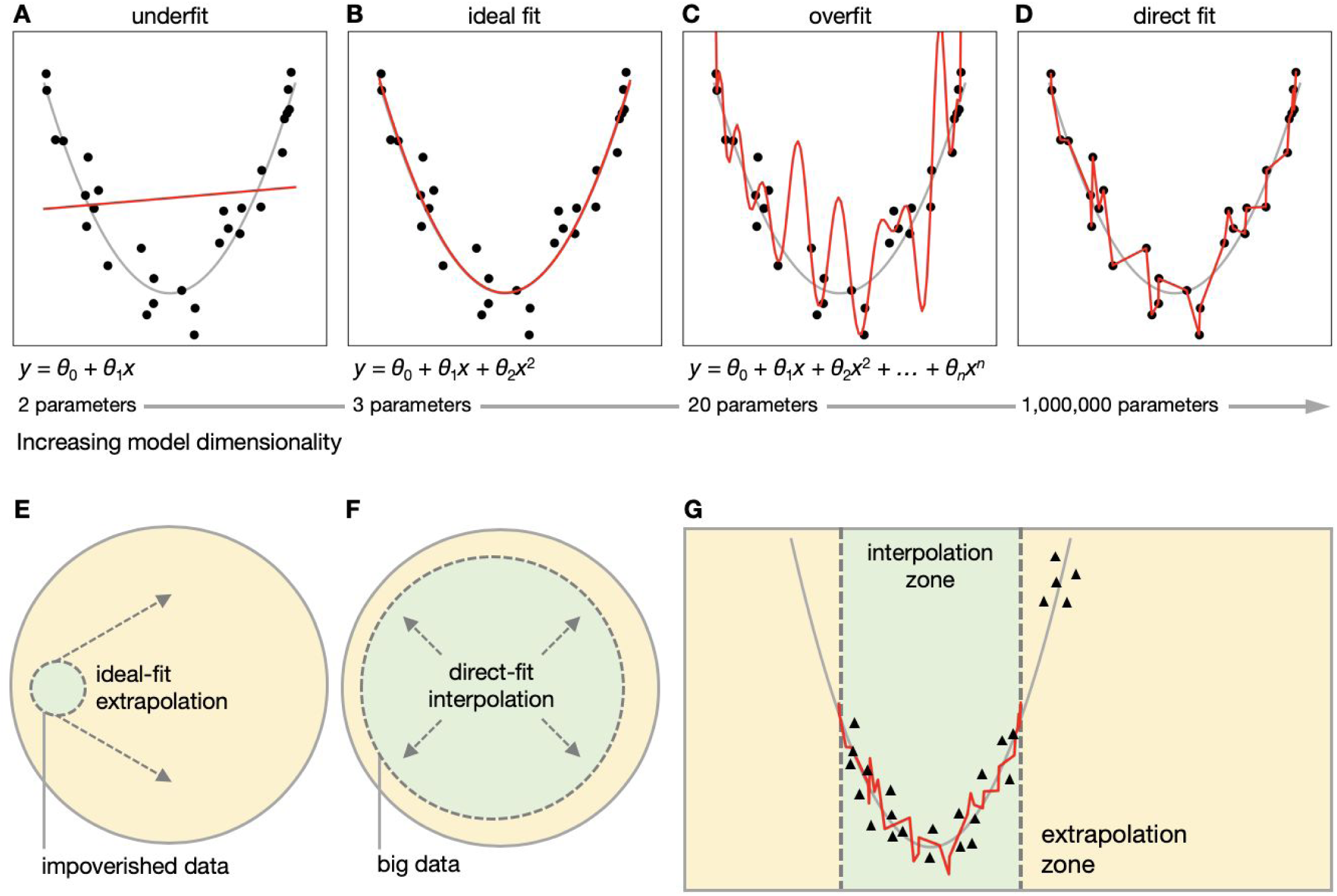
Direct-fit learning with dense sampling supports interpolation-based generalization. (***A***) An overly simplistic model will fail to fit the data. (***B***) The ideal-fit model will yield a good fit with few parameters in the context of data relying on a relatively simple generative process; in fact, this is the model used to generate the synthetic data (with noise) shown here. (***C***) An overly complex (i.e., over-parameterized) model may fixate on noise and yield an explosive overfit. Panels *A*–*C* capture the “textbook” description of underfitting and overfitting. (***D***) Complex models such as ANNs, however, can nonetheless yield a fit that both captures the training data and generalizes well to novel data within the scope of the training sample (see Panel G and Bansal et al., 2018, for a related discussion). (***E***) Traditional experimentalists typically use highly-controlled data to construct rule-based, ideal-fit models with the hope that such models will generalize beyond the scope of the training set, into the extrapolation zone (real-life data). (***F***) Direct-fit models—like ANNs and, we argue, BNNs—rely on dense sampling to generalize using simple interpolation. Dense, exhaustive sampling of real-life events (which the field colloquially refers to as “big data”) effectively expands the interpolation zone so as to mimic idealized extrapolation. (***G***) A direct-fit model will generalize well to novel examples (black triangles) in the interpolation zone, but will not generalize well in the extrapolation zone.

We contend that this textbook view should be revised to account for the fact that, in a data-rich setting, over-parameterized models can provide a mindless, yet powerful, form of generalization. Any model is designed to solve a particular type of problem, and the problem to be solved changes drastically when we shift from preferentially sampling a limited parameter space in a controlled experimental setting to densely sampling a wide parameter space using big data in a performance-oriented real-life setting.

#### Generalization based on impoverished data

When the scope of the data is narrow relative to the scope of world’s possible states (Figure 1E), over-parameterized models will tend to learn idiosyncrasies specific to the training data and will not extrapolate beyond that scope. This well-curated, narrow sampling aperture is what we have in mind when we teach introductory statistics using diagrams like Figure 1A–C. For example, only the ideal-fit model revealing the underlying generative parabola rule (*y* = *θ*_0_ + *θ*_1_*x* + *θ*_2_*x*^2^) can be useful for predicting the values of new observations in the extrapolation zone in Figure 1E. In contrast, the underfit and overfit models will be useless in predicting the values of any new point in the extrapolation zone. In other words, such generative ideal-fit models provide the ultimate model for generalization, which relies on a complete understanding of the underlying rules used to generate the observations. However, extrapolation-based generalization requires that the generative rules hold outside of the training zone (e.g., simulated data). In cases where there are complex nonlinearities and interactions among variables at different parts of the parameter space, extrapolation from such limited data is bound to fail. (The validity of this assumption about the uniformity of parameter space is difficult to empirically evaluate and may vary wildly across domains of inquiry; here we sample from a simple distribution for the purpose of simulation, but the world around us clearly does not resemble such a simple generative process.)

The perspective of the narrow aperture (Figure 1E), from which we can uncover the underlying generative rules needed to predict observations in a wide variety of contexts based on data collected during contrived and highly-controlled experiments has a privileged role in the minds of scientists across many disciplines, including physics, chemistry, neuroscience, and psychology. Interestingly, many computational neuroscientists, cognitive and developmental psychologists, and psycholinguists adopt this narrow aperture image when theorizing about the neural code. This creates a tension: experimentalists use contrived stimuli and designs to recover elegant coding principles (e.g., Hubel and Wiesel, 1962); but it remains unclear whether these principles actually capture neural responses in naturalistic contexts (Felsen and Dan, 2005; Olshausen and Field, 2005; Hasson and Honey, 2012; Hamilton and Huth, 2018). This is not a flaw of experimental design per se; cleverly-designed experiments can in fact expose principles of direct fit. However, the limited generalizability of experiments using contrived, non-representative manipulations is often glossed over (Brunswik, 1947). Historically these practices and tensions can in part be traced to an argument from cognitive psychology that the brain is not exposed to rich enough data from the environment to navigate the problem space (Chomsky, 1965). Therefore, to predict novel outcomes in novel contexts, the neural code is assumed to rely on implicit generative rules (either learned or inherent).

#### Generalization based on big data

Dense sampling of the problem space (Figure 1F) can flip the problem of prediction on its head, turning an extrapolation-based problem into an interpolation-based problem. This is illustrated in Figure 1G, when we add new observations (black triangles) not seen during training to the interpolation (green) zone. Counterintuitively, within the interpolation zone, over-parametrized models with sufficient regularization (Figure 1D), which we denote as direct-fit models (see section below), can attain as good predictive performance as the ideal-fit model (if not better, under conditions in which variability in the data is not due to random noise).

Interpolation is a local process, which does not rely on explicit modeling of the overarching generative principles. It uses simple, local heuristics, like nearest neighbors or averaging, to place the current observation within the context of past observations. Furthermore, as will be discussed below, over-parametrized models provide new computational tools to learn complex multidimensional statistical regularities in big data, where no obvious generative structure exists.

To summarize this point: interpolation uses local computations to situate novel observations within the context of past observations; it does not rely on explicit modeling of the overarching generative principles. Unlike extrapolation, interpolation was thought to provide a weak form of generalization because it can only predict new data points within the context of past observations. Thus, when we considered the brain, we have traditionally assumed that interpolation did not provide a sufficient form of generalization to support complex behavior, as the task of the brain is to extrapolate from a small number of examples to a near-infinite range of possible observations (Figure 1E). But this problem only arises if the scope of the training space is small or impoverished (as in highly-controlled experiments). However, the move to big data reframes the problem (Figure 1F): if we densely sample parameter space using millions of free parameters to robustly fit millions of examples, there is remarkable power in simple interpolation-based predictions (see Box 2).

### Direct fit and artificial neural networks

Not all over-parameterized models overfit the data. There are two types of over-parameterized models: explosive overfit and direct-fit. In the case of explosive overfit (Figure 1C), the model memorizes all training data points, but otherwise strays wildly from the underlying structure of the data and does not afford interpolation or extrapolation. The direct-fit model also relies on over-parameterization to match the data structure. In contrast to explosive overfit model, however, the direct-fit model regularizes the process to avoid explosive overfit while optimizing the alignment to the structure of the training data (Figure 1D). This regularization may collapse redundancies, imposing priors for sparseness or smoothness, but, critically, can be implemented using generic, local computations, and does not require any explicit model of the latent features of the data.

As an example of a direct-fit procedure, we will use standard ANN architectures to model two low-dimensional processes. For a brief discussion of ANNs and their relation to BNNs, see Box 1. We will use two architectures: a standard fully-connected ANN for testing interpolation and extrapolation over space, and a recurrent neural network for testing interpolation and extrapolation over time.

#### Box 1: Artificial and biological neural networks

Artificial neural networks (ANNs) are formal learning models inspired by the biological neural networks (BNNs) that constitute living brains. ANNs, however, are an extreme abstraction of BNNs. Typically, biological neurons have three main structures: the cell body, axon, and dendrites. They come in a variety of shapes and functions, classified into unipolar, bipolar, and multipolar groups, each further subdivided into a menagerie of different types. Each neuron in a BNN is modulated by a specific set of neurotransmitters and embedded in a complex local neuronal circuit with different input and output units, inhibitory lateral connections, and a unique layout of interconnectivity. In addition to varying local circuit architecture, some biological nervous systems include functionally specialized systems-level components, like the hippocampus, striatum, thalamus, and hypothalamus, which are not generally included in current ANNs. Furthermore, the dynamic and biophysical properties of biological neural networks are vastly different from ANNs. Finally, most ANNs are disembodied and do not interact closely with the environment in a closed-loop fashion (see Box 3). This degree of abstraction has led many neuroscientists to dismiss ANNs as irrelevant for understanding biological brains.

### Generalization of ANNs in the interpolation and extrapolation zones

To illustrate the properties of direct-fit models, we first trained an ANN on a set of 10,000 training examples of even numbers (green dots) sampled with variance from a simple sine function (Figure 2). The ANN was trained to predict the *y*-axis values from the *x*-axis values (imitating a spatial task). The ANN was composed of one input neuron, three fully connected hidden layers, each with 300 neurons, and one output neuron. Even such a small network of 902 neurons results in an over-parameterized model with approximately 180,600 adjustable parameters (weights). The model was trained with simple backpropagation through stochastic gradient descent.

**Figure 2.**
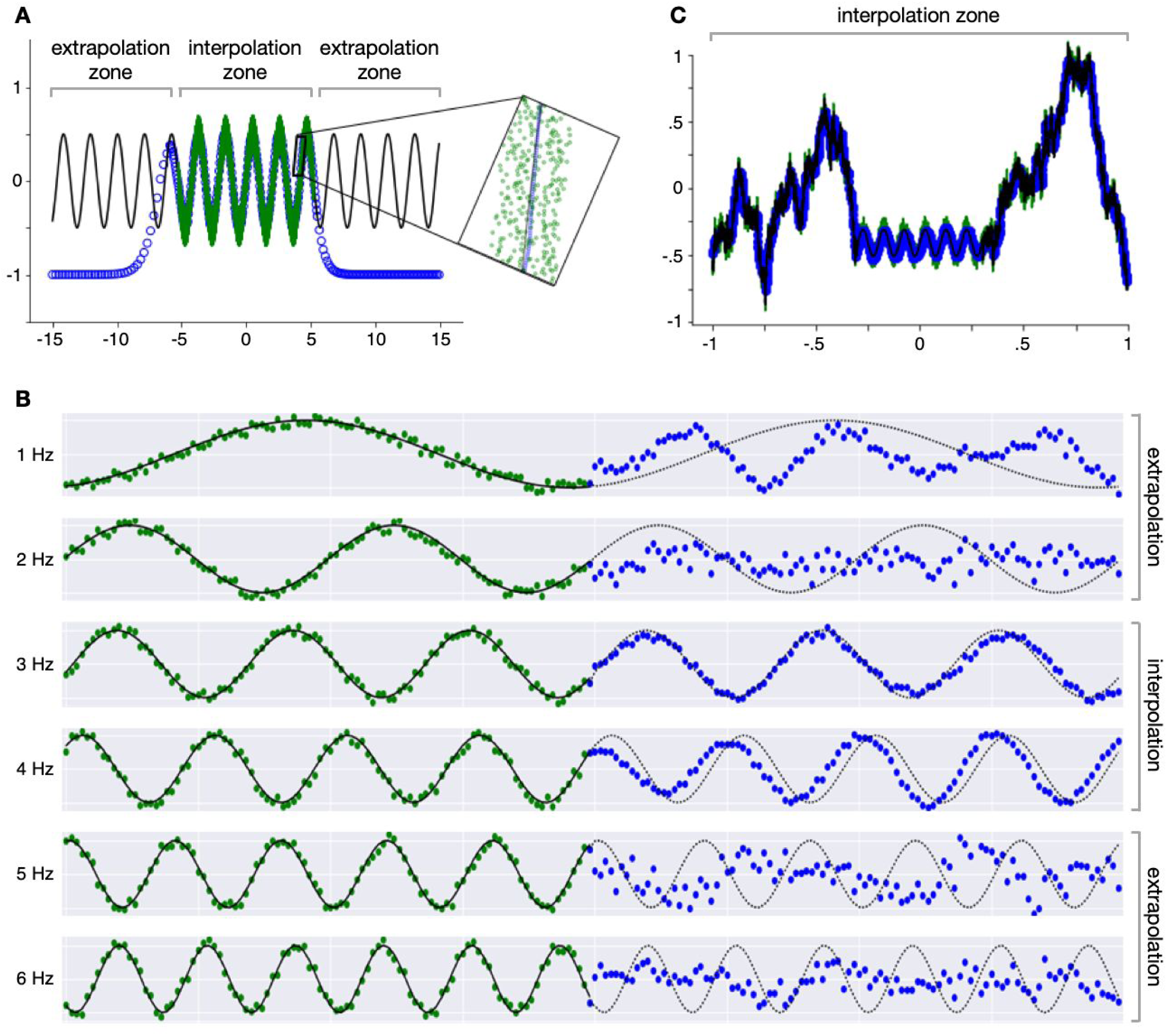
ANNs can only generalize within the interpolation zone. (***A***) Interpolation over space: A simple ANN model with three fully connected hidden layers was trained to predict the output of sine function mapping *x*-axis to *y*-axis values. Training examples (green markers) were *x* values between −5 and 5 (comprising only even values). Predictions for test *x* values ranging from −15 to 15 (comprising only odd values) are indicated using blue markers. The ideal sine wave (from which the observations are sampled) is indicated by the black line. The model was able to generalize to new test examples not seen during training within the interpolation zone, but not within the extrapolation zone. (***B***) Interpolation over time: A simple recurrent ANN (LSTM) was trained to predict the sequence of forthcoming observations from a sine function. Training examples were sampled from the first half of sine wave sequences between 2.5 and 4.5 Hz. The trained model was supplied with test samples from the first half of a sequence (green markers) and predicted the subsequent values (blue markers). The model was able to generalize to new frequencies not seen during training within the interpolation zone, but not within the extrapolation zone. (***C***) Interpolation provides robust generalization in a complex world: Given a rich enough training set, the advantage of direct-fit interpolation-based learning becomes apparent, as the same ANN from panel A is able to learn an arbitrarily complex function (for which there is no ideal model).

All training examples were sampled from a confined parameter space (−5 < *x* < 5), which we denote as the interpolation zone. After training, the model was used to predict the *y* value for 10,000 new examples (even *x* values; blue dots) sampled at a wider range of values (−15 < *x* < 15) extending beyond the interpolation zone into the extrapolation zone. Our goal was to measure the ability of the direct-fit model to interpolate and extrapolate the values of the new test examples not seen during the fitting process.

By construction, an ideal sine function (black line, Figure 2A), a model with exactly one free parameter, will achieve optimal prediction of all blue points in the interpolation and extrapolation zones. The ANN, however, managed to predict new observations not seen during training (Figure 2A) only within the interpolation zone. The ability of the direct-fit model to interpolate but not to extrapolate is clearly seen when we look at the test data points in Figure 2A. The direct-fit model does not produce any clear rule for how the data should look like outside the context or “scope” of the interpolation zone, providing a poor prediction for new examples in the extrapolation zone. However, within the interpolation zone, the ANN is as good as the ideal-fit model in predicting the values of new observations not seen during training. This can be seen in the magnified portion of Figure 2A. Note how the predicted values (blue points) overlap with the sine function used to generate the data (black line). The interpolation zone is closely related—but not identical—to the training set. The interpolation zone corresponds to the region of parameter space spanned by the training samples, but can contain an infinite number of novel samples not observed during training.

While in this case, the ANN did not truly learn the ideal sine function necessary for extrapolation, it was still capable of optimizing the fit to achieve high prediction quality within the interpolation zone. One could argue that the ANN has implicitly “learned” the sine function within the interpolation zone, but the critical distinction is that this implicit representation of the sine function is an incidental or emergent byproduct of the structure of the input and the fitting procedure. We can interrogate the ANN for representations resembling the sine function, but these exist only because we injected them into the training data; the ANN has simply learned how to interpolate new observations within the scope of the training set. By analogy, it may be misleading to claim that the brain represents in any fundamental way some experimental variable, even in experiments where such a description can account for a considerable amount of variance in the neural responses (Marom et al., 2009).

We demonstrate similar behavior, but now over time rather than space, when training a recurrent long short-term memory neural network (LSTM; Hochreiter and Schmidhuber, 1997) to learn sine wave sequences (Figure 2B). In this case, instead of using a fully-connected ANN to learn the spatial relationship between *x* and *y* values, we trained the LSTM to predict a future sequence of *y* values based on the preceding sequence of 100 *y* values sampled within a 1-second input window (green). The network was trained on sine functions cycling at different frequencies from 2.5 to 4.5 Hz (training zone; excluding samples at exactly 3 and 4 Hz). To assess the network’s capacity for interpolation and extrapolation, we tasked the trained network with predicting the values of a forthcoming sequence of 100 *y* values at novel frequencies not sampled during training, either within the interpolation zone (i.e., 2.5–4.5 Hz) or in the extrapolation zone (i.e., frequencies slower than 2.5 Hz or faster than 4.5 Hz). The LSTM was able to predict the next 100 *y* values for new sine waves not seen during training, but only at frequencies within the interpolation zone (e.g., at 3 and 4 Hz in Figure 2B). The LSTM failed to extrapolate in predicting values for new sequences at frequencies outside the interpolation zone (i.e., 1, 2, 5, and 6 Hz in Figure 2B).

The “no free lunch” theorem demonstrates that optimization for one task will necessarily deteriorate performance in another (Wolpert and Macready, 1997). Here, we see how introducing a different architecture can improve prediction of a sine function at a particular frequency. However, this will not solve the extrapolation problem in general, as the network still does not learn the ideal, rule-based sine function required to extrapolate to all sine waves, but simply learns how to interpolate new observations within the scope of the training set. While increasingly sophisticated models trained on rich data may eventually approximate the human brain’s exceptional robustness to broadly-distributed spatial and temporal structures, both ANNs and BNNs are nevertheless subject to the no free lunch theorem. They learn ad hoc solutions by optimizing for a narrow region of problem space, and a single architecture cannot excel in every domain (Gomez-Marin and Ghazanfar, 2019). In the same vein, evolution yields organisms that are optimized to fit the constraints of a given ecological niche (e.g., the deep sea or the desert), but does not find “well-designed” or globally optimal solutions to survive everywhere on the globe.

### The robustness of direct-fit

The ideal sine function allows us to extrapolate to infinite new values. In contrast, the over-parameterized direct-fit model can only be used to predict values of new observations within the confined interpolation zone. Here, we have artificially limited the underlying structure of the data such that the process generating observations can be captured in one parameter. To draw an analogy with cognitive psychology, we have constrained the experimental design to parametrically vary a single stimulus feature (e.g., the spatial frequency of a Gabor filter), holding all other environmental variables constant. We have started with a simple model to generate observations in hopes of recovering the original generative rule from which the training data were sampled. In fact, when interrogating the over-parameterized model under these conditions, we simply recovered the task dimensions by which we constructed the experimental paradigm or training set (Gao et al., 2017; Stringer et al., 2019).

In contrast to ideal-fit models, which flourish in simulations and well-defined experimental settings, direct-fit models can provide powerful ways to model big data in which the latent structure is multidimensional, complicated, and prohibitively difficult to model using a handful of factors. For example, consider a world (Figure 2C) in which the underlying sine function only applies to a narrow range of training examples (between −5 > *x* > 5), but beyond that specific range the sine function no longer describes the data structure. That is, when the data is sampled over a wider range of training examples (between −15 > *x* > 15), it behaves in a consistent and stable manner, which is, however, very different from the sine wave (to drive the point home, we generated these samples using a simple random-walk algorithm, which by construction generates an arbitrary function).

As in Figure 2A, we retrained the same over-parameterized ANN to fit 30,000 even-valued observations (green points) sampled from a wider parameter space (between −15 > *x* > 15). Due to its flexibility and adaptivity, the over-parameterized ANN model can now interpolate to accurately predict the values of 30,000 new observations (blue points) not seen by the model within the wider training zone. Note, in contrast to the ideal-fit model, the direct-fit model does not catastrophically fail at this wider range of training examples—the model is expressive enough to fit whatever stable data structure it observes. Indeed, as presented schematically in Figure 1E–F, direct-fit models thrive in the context of big data, where the interpolation zone increases with the scope of the training set.

By widening the interpolation zone, the model’s inability to extrapolate becomes less and less of a liability (Feldman, 2019). The same direct-fit procedures can be expanded to fit arbitrarily complex data structures (Cybenko, 1989; Funahashi, 1989; Hornik et al., 1989; Raghu et al., 2017). The ability of over-parameterized models to robustly fit complex data structures provides unparalleled predictive power within the interpolation zone, making them uniquely suitable for multidimensional, real-life situations for which no simple, ideal model exists. Ultimately, as we develop new architectures and learning rules, we predict that these models will only be limited by the scope of their training observations and the complexity of the task (Figure 1F). In other words, when the data structure is complex and multidimensional, a “mindless” direct-fit model, capable of interpolation-based prediction within a real-world parameter space, is preferable to a traditional ideal-fit explicit model that fails to explain much variance in the data.

### What is needed for successful direct fit?

Over-parameterized models are notorious for being hyper-expressive, prone to imposing imaginary structure on random unstructured training sets. For example, it was shown (Zhang et al., 2016) that ANNs can be trained to fully memorize arbitrary associations between a set of object labels and a set of randomly shuffled images that do not match the labels. In this case, the network memorized the entire arbitrary training set, achieving close to 100% classification accuracy on the training data—but with no generalization to a new unseen set of test images (i.e., poor interpolation). The exact same set of images and labels were then used to train the same deep network, but this time the images were matched with correct labels. Similar to the random labels condition, the network achieved close to 100% classification on the training set, but in this case, the model did not overfit; rather, it was capable of generalizing and correctly labeling new test images not seen during training.

What is the difference between these two cases that relied on the exact same stimuli, network architecture, learning rules, and objective function, but resulted in such different models? The solution to this puzzle lies not in the features of the model, but rather in the properties of the external world. There are five requirements for over-parameterized models to generalize: (1) they must be fit to a structured world; (2) the world must be sampled densely and widely; (3) the model must support a high-dimensional encoding space; (4) the model must have the correct objective function(s); and (5) the model must implement effective regularization during optimization to avoid explosive overfit.

#### The structure of the world

The world is hardly random—it is structured according to laws of physics, biology, sociology—and the mind reflects this structure. However, unlike ideal-fit models, the nervous system does not explicitly define some handful of relevant signal dimensions. An over-parameterized direct-fit model with sufficient sampling is flexible enough to integrate multidimensional signals for interpolation. For an illustrative example, consider the faces of people around you. We carry our faces with us everywhere we go, and although we slowly age, we retain enough features over time for people to recognize us at around 97% accuracy across different situations and across time (O’Toole et al., 2018). When the signals are unstable, however, direct-fit models are likely to fail. For example, in a world in which we sporadically swap facial features, or in which we share identical facial features with all other people, the task of face recognition would be much more difficult. Drastic, qualitative deviations from the structure of our familiar world would likely result in a catastrophic failure in interpolation-based generalization, but we hope to rarely, if ever, encounter situations that would require such extrapolation (the impending climate collapse notwithstanding).

#### Dense sampling of the world

In real life, sensory signals are usually noisy and dynamic. For example, although our facial features are relatively stable, we may look very different under different lighting conditions, from different angles, with different make-up and hairstyles, or when occluded by different objects. For direct-fit to work, we need to densely sample a broad parameter space (Figure 1F) to ensure robust interpolation. For example, if we were to fit a model to only forward-facing face images, generalization to profiles would be poor because profile images fall outside the interpolation zone along the dimension(s) of viewpoint (Srivastava and Grill-Spector, 2019). If, however, we were to sufficiently sample images across different viewpoints, lighting conditions, and different states of occlusion, we would be able to interpolate across all these dimensions. Similarly, if we were to train a model only on images of one face, it wouldn’t be able to recognize anyone else in the world. If we were to train the model on millions of Western faces, it would likely recognize Western faces, but extrapolate poorly to East Asian faces (Malpass and Kravitz, 1969; O’Toole et al., 2018). From this perspective, the brain is not necessarily an expert in face recognition per se, but rather it is expert in recognizing the faces it generally encounters (Ramon and Gobbini, 2018; Young and Burton, 2018). That is, our face recognition behavior does not necessarily imply that our brain learns an ideal, low-dimensional model of faces that it can use to extrapolate to new, unfamiliar faces. Rather, we densely sample face space over a range of parameter values broad enough to roughly circumscribe most of the faces we encounter, thus enabling interpolation (see Box 2 for details).

##### Box 2. Face recognition and language models: two examples of direct fit

We argue that BNNs and ANNs belong to the same family of direct-fit optimization models. Nonetheless, across different biological and artificial networks, there is considerable variability in circuit architecture, learning rules, and objective functions. Although novel computational motifs regularly emerge from the machine learning literature, the space of possible models is vast and largely unexplored.

#### High-dimensional encoding space

For direct-fit to work, we need to adjust millions of parameters to accommodate the complex, multidimensional structure of the world. In ANNs, these parameters correspond to the synaptic weights between numerous simple computing elements. In practice, this high-dimensional multivariate encoding space typically captures the structure of the world in distributed embeddings. Any feature of the world is represented across many computing elements and each computing element participates in encoding many features of the world. This distributed encoding scheme has several useful properties, including high capacity/expressivity, robustness to noise (e.g., graceful degradation), and, critically, approximate continuity in vector space that natively supports interpolation-based generalization (Hinton et al., 1986). On the other hand, this encoding scheme makes it difficult to interpret the functional tuning of any single unit or neuron (e.g., Ponce et al., 2019). Modern ANNs have exposed the power and versatility of this encoding scheme: a variety of seemingly distinct “tasks” can be performed by interpolating over a single high-dimensional embedding space (e.g., Eliasmith et al., 2012; O’Toole et al., 2018; Radford et al., 2019; Raffel et al., 2019).

#### Ecological objective functions

Over-parameterized models are often hyper-expressive, and can fit essentially any dimension of the data or world. However, most dimensions are likely to contain little, if any, functional advantage for the organism. Objective functions drive optimization of the model weights to fit to the desired dimensions (Marblestone et al., 2016). There are two types of objective functions, internally guided (which are sometimes referred to as unsupervised, but we prefer the term of “self-supervised”) and externally guided (referred to as supervised, but we prefer the term of “externally-supervised”). Only a small set of objective functions will yield models supporting adaptive behavior, and such objectives may propagate across brains and across generations (spreading even faster among social organisms). On the other hand, uninformative objective functions may be useless or costly, and overall less rewarding. For example, a training set of 10,000 face images can be divided to 2^10,000^ groups, but only a subset of these subdivisions are functionally meaningful. Examples of useful subdivisions may include gender, identity, or age. Examples of less useful subdivisions may include hairstyle, eye color, shape of the nose, length of eyelashes, or the number/location of beauty marks, blisters, freckles, and so forth. Most ANNs could be trained to prioritize any of these features and perform remarkably well were we to assign the network such an objective function (Marblestone et al., 2016). By allowing the system to converge on functional solutions, while remaining largely blind to the global, underlying structure of the world, adaptive objective functions in learning are closely related to selection pressures in biology, as discussed below.

#### Effective regularizations procedures

Regularization effectively imposes a prior on optimization processes to prevent explosive overfitting. Again, we can draw on the analogy to evolution, where the predominantly incremental nature of genetic variation, robustness to genetic mutations, and constraints of physiology (imposed both morphologically and due to limited resources) regularize the fitting process. In fact, the genome may impose exceptionally strong priors on learning (Zador, 2019).

### The black box argument

When applied to suitable data using the appropriate objective functions, direct-fit optimization procedures can provide us with powerful functional models that use interpolation to predict the values of new observations in real-world contexts. As demonstrated in Figure 2, these models do not explicitly encode the generative structure of the data and lack the ability to extrapolate to previously unseen contexts.

Critics often refer to over-parameterized direct-fit models pejoratively as “black-box” models: models that, given the correct input, generate the correct output, without any explanation of their internal workings (Ashby, 1956; McCloskey, 1991). For example, the human face network is comprised of millions of neurons and billions of synaptic weights, which as an ensemble are capable of recognizing faces of thousands of individuals across different views and contexts (Jenkins et al., 2018). Similarly, using deep neural networks, and without hardwiring or even endeavoring to “explain” the latent facial features or rules by which their models perform, commercial face-recognition software can recognize faces with (super)human accuracy (Taigman et al., 2014; Schroff et al., 2015). Thus, one may argue, such ANNs have simply duplicated the original problem by creating one more black box model for face recognition; as if the brain wasn’t enough.

We argue that there is nothing opaque about ANNs—they are fully transparent “glass boxes”. The physicist Richard Feynman famously wrote on his blackboard, “What I cannot create, I do not understand.” We build artificial networks according to explicit architectural specifications; we train networks using explicit learning rules and finite training samples with well-specified objective functions; we have direct access to each weight in the network. Given their unprecedented level of transparency, why do we deem ANNs black box models? We do so because we are deeply committed to the assumption that the ANN must learn a set of human-interpretable rules necessary for processing information. This is our classical criterion for understanding. Since we do not readily find such rules when interrogating the distribution of millions of adjustable weights within over-parameterized artificial (and biological) neural networks, we demote such models to black box status (Lillicrap et al., 2019).

In contrast to the common black-box argument, which fixates on the interpretability of the fitted model parameters, we argue that the broad family of direct-fit neural network models actually provide a concise framework for understanding the neural code. Artificial neural networks can be understood in terms of three components: network architectures, learning rules, and objective functions (Richards et al., 2019). Although BNNs differ substantially from ANNs in all three factors (see Box 1 and 3), both belong to the same family of direct-fit models. BNNs, however, are the result of billions of years of evolution in a complex world, while ANNs are in their infancy. Nonetheless, ANNs provide a proof of concept that neural machinery may rely on mindless fitting over exhaustive samples to enable powerful interpolation-based generalization performance. There is a surprising simplicity in the design specifications of direct-fit ANNs and BNNs, but this simplicity does not guarantee the interpretability we initially sought.

Direct-fit models do not learn rules for extrapolation, but rather use local interpolations to determine the value of new examples based on their proximity to past examples within a multidimensional embedding space (see Box 2). BNNs and ANNs, from this perspective, belong to a family of weakly representational models, capable of learning the mapping between input and output using direct-fit optimization procedures, while being effectively agnostic as to the underlying structure of the world. We should exercise caution in cases where these models seem to “learn” simple, psychologically interpretable variables. It can be tempting to impose our own intuitive or folk-psychological interpretations onto the fitted model, but this is misguided. If a generic network learns such a rule, this rule is likely inherent in the training set, and is thus not so much a meaningful property of the network as it is a property of the data (see Figure 2). These interpretable rules arise incidentally, as an emergent byproduct of the fitting procedure. The incidental emergence of such rules is not a “goal” of the network and the network does not “use” the rules to extrapolate. This mindset, in fact, resembles pre-Darwinian teleological thinking and “just-so stories” in biology (Gould and Lewontin, 1979; Mayr, 1992). Evolution provides perhaps the most ubiquitous and well-known example of a biological fitting process that learns to act in the world while being blind to the underlying structure of the problems or their optimal solutions.

### The power of adaptive fit in evolution

Most biological processes are not guided by the explicit objective of understanding the underlying structure of the world. Evolutionary theory aims to explain how complex organisms (ranging from amoebae, to plants, fungi, fish, and mammals) and complex biological mechanisms (such as photosynthesis, gills, wings, retinas) evolved to fit their local ecological niches, without any explicit comprehension of the problems at hand, and without any understanding of the solutions to overcome them (Darwin, 1859). Evolution is the study of ever-changing, blind, local processes, by which species change over time to fit their shifting local environment (Fisher, 1930; Williams, 1966).

The theory of evolution tries to explain the blind, local fitting processes by which all living creatures on earth have evolved (Figure 3). These organisms all share the same origin and their evolution relies on a handful of basic processes (Lewontin, 1970; Gould, 1982): (*a*) *over-production with variation* via genetic mutation, gene regulation and expression, genetic drift, endosymbiosis, or hybridization; (*b*) *inheritance* via vertical transmission of genetic material from parent to offspring, and horizontal transmission of genetic material between unicellular and/or multicellular organisms; (*c*) *combinatorial power* of the genetic code to support diverse morphologies and organismal complexity; (*d*) *selection* via natural and artifactual external forces, sexual, kin, and group preferences; and (*e*) *time* necessary to support the iterative diversification and refinement of the phylogenetic tree, which has been unfolding incrementally over many generations for over 3.5 billion years.

**Figure 3.**
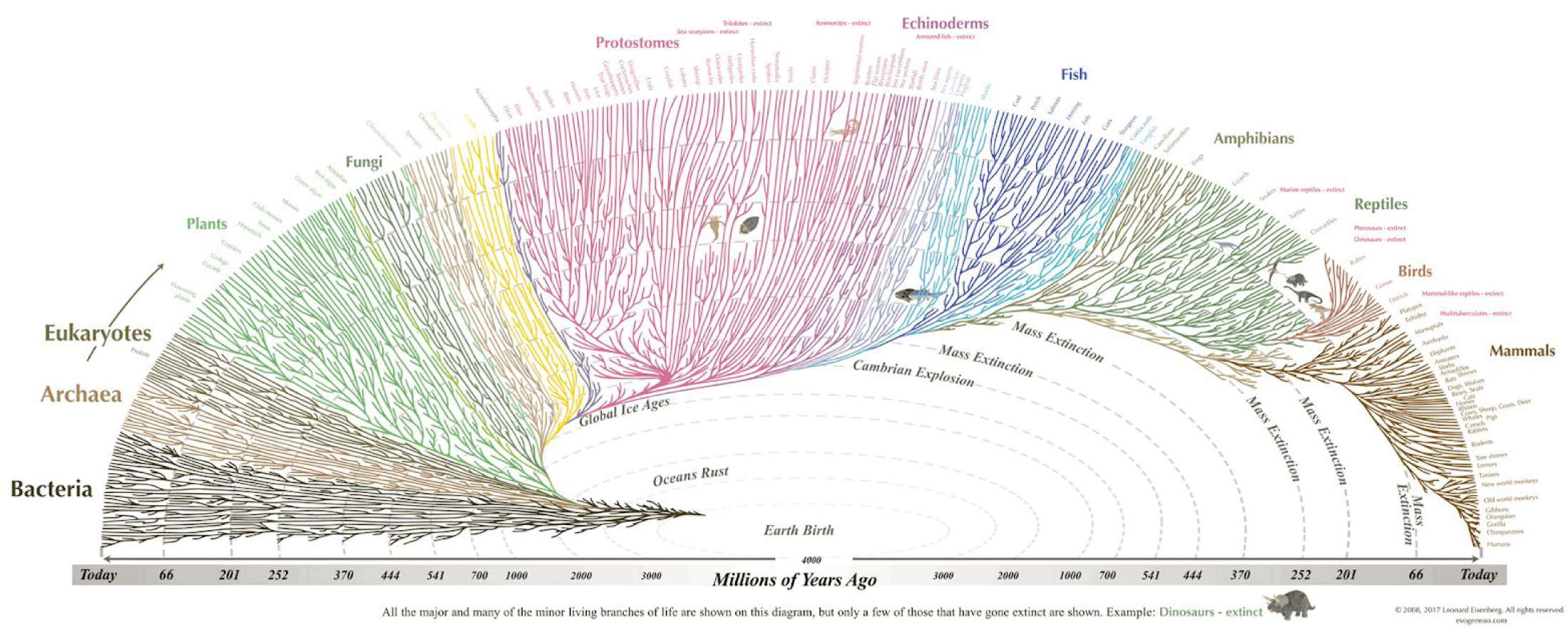
Evolution by natural selection is a mindless optimization process by which organisms are adapted over many generations according to environmental constraints (i.e., an ecological niche). This artistic rendition of the phylogenetic tree highlights how all living organisms on earth can be traced back to the same ancestral organisms. Humans and other mammals descend from shrew-like mammals that lived more than 150 million years ago; mammals, birds, reptiles, amphibians, and fish share a common ancestor—aquatic worms that lived 600 million years ago; and all plants and animals derive from bacteria-like microorganisms that originated more than 3 billion years ago. Reproduced with permission from Leonard Eisenberg (https://www.evogeneao.com).

The theory of evolution makes use of a few simple principles to explain the tight connections between vast arrays of phenomena. Thus, the theory of evolution is simple and parsimonious. At the same time, evolution is inefficient and costly in its implementation, given that today’s organisms have evolved over billions of years of local interpolations (Thompson, 2013). Moreover, in contrast to the laws of nature in physics, which provide us with the ability to extrapolate and predict events in different corners of the universe, evolution is a local process, not easily used for extrapolation to the next evolutionary step. Predicting the forthcoming ramifications of the tree of life on Earth one million years from now is prohibitively difficult. Similarly, we cannot easily predict the morphology of an organism given a novel set of environmental constraints; that is, the theory of evolution cannot be used to extrapolate phylogenetic trees beyond planet Earth, in ecological niches at different corners of the universe. Does the lack of extrapolation undermine the explanatory power of the theory of evolution? Should we admit that we simply do not understand evolution because the fitting procedure does not yield a finite set of intuitive, psychologically interpretable biological motifs and organisms?

### Direct-fit to nature

The critical and subversive advance of evolutionary theory was to remove the need for an “intelligent” force to guide change (Dawkins, 1986; Dennett, 1995). Similarly, direct-fit neural networks remove the need for intentional or interpretable rules to guide learning (Dennett, 2017; Table 1). The ANN does not require the engineer to inject into the network human-interpretable rules describing, e.g., face configuration, nor should the engineer impose these interpretations on the network’s solution. Evolution teaches us how endless iterations of the same blind process of variation guided by natural selection can produce the rich variety of organisms and biological mechanisms we observe in nature. Similar to natural selection, the family of models to which both ANNs and BNNs belong optimize parameters according to objective functions to blindly fit the task-relevant structure of the world, without explicitly aiming to learn its underlying generative structure. In fact, evolutionary algorithms often find non-intuitive solutions to complex problems, especially in the context of multiple overlapping or conflicting objectives (Holland, 1975; Bäck, 1996; Eiben and Smith, 2015). An organism’s genome, analogous to a given ANN architecture, implicitly encodes certain structural assumptions about the statistics of the world and objective functions (Maynard Smith, 2000; Godfrey-Smith, 2007; Adami, 2012; Zador, 2019). Both genome and neural network are highly expressive, distributed encoding architectures (Quackenbush, 2001; Raghu et al., 2017). In this sense, network solutions adapted to performing particular behaviors are analogous to organisms adapted to particular niches as guided by evolution. In the same way that ANNs fail at extrapolation, an organism transplanted outside the ecological niche to which its species has adapted may perish.

**Table 1.**
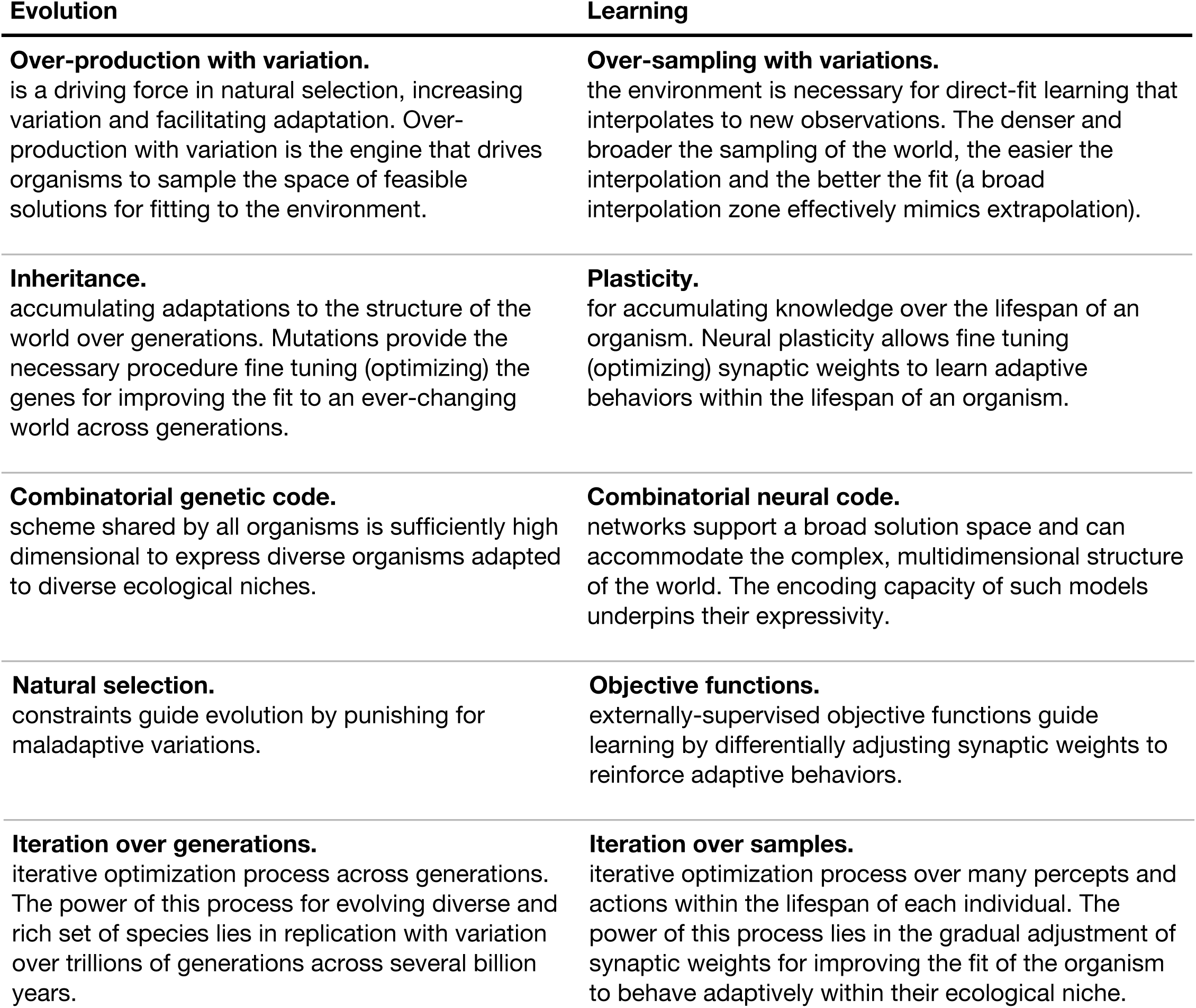
The parallels between direct-fit learning and evolution by natural selection.

Evolution does not have the luxury of operating in an idealized, highly-controlled parameter space (like an experimenter’s laboratory), and neither do biological learning organisms (Anderson and Chemero, 2016). Therefore, much like optimization in deep learning, evolution by natural selection puts a premium on behavior and task performance; interpretability in the phenotypes it yields is only happy coincidence.

### Direct-fit models contradict three basic assumptions in cognitive psychology

From its inception, cognitive science has argued against over-parameterized direct-fit models, asserting that cognition materializes under three fundamental constraints. First, the brain’s computational resources are limited, and the underlying neural code must be optimized for particular functions (e.g., Chomsky, 1980; Fodor, 1983). Second, the brain’s inputs are ambiguous and too impoverished for learning without built-in knowledge (e.g., Chomsky, 1980). Lastly, shallow externally supervised and self-supervised methods are not sufficient for learning (e.g., Pinker, 1994). Briefly, in the example of grammar learning, both the linguistic input and feedback are claimed to be insufficient; therefore, language learning must rely on hardwired (i.e., not learned) computational modules to support our generative capacity to extrapolate (Chomsky, 1965; cf. Pullum and Scholz, 2002; Ramscar and Yarlett, 2007; Christiansen and Chater, 2008). Considering the brain as a biological neural network using direct-fit optimization challenges these three assumptions and proposes new routes for learning.

#### Computational resources are not scarce

Each cubic millimeter of cortex contains hundreds of thousands of neurons with millions of adjustable synaptic weights, and biological neural networks utilize complex circuit motifs hierarchically organized across many poorly understood cortical areas (Felleman and Van Essen 1991). Thus, relative to BNNs, ANNs are simplistic and minuscule. Relative to the ideal-fit models, however, the sheer size of ANNs with millions of parameters and biological networks with billions of parameters seems overwhelming. Although the brain is certainly subject to wiring and metabolic constraints, we should not commit to an argument for scarcity of computational resources as long as we poorly understand the computational machinery in question (Levy et al., 2004).

While the capacity to learn simple tasks from big data may be practically unbounded given the expressivity of ANNs, other architectural constraints may impose structural constraints on the capacity of the system to learn and act in the world (either chew or talk with your mouth). Such constraints may include the need to integrate information across modalities and timescales, while selecting and executing a small set of coherent behaviors at each moment (Musslick et al., 2017).

#### The input is not impoverished

Direct-fit relies on dense and broad sampling of the parameter space for gaining reliable interpolations. One of our main insights is that dense sampling changes the nature of the problem and exposes the power of direct-fit interpolation-based learning (Figs. 1 and 2). Quantifying the input entering the brain is a complicated and laborious task (Sivak, 1996). Recent measurements suggest that the incoming input may be vast and rich (Zyzik, 2009). For example, we may be exposed to thousands of visual exemplars of many daily categories a year, and each category may be sampled at thousands of views in each encounter, resulting in a rich training set for the visual system. Similarly, with regard to language, studies estimate that a child is exposed to several million words per year (Roy et al., 2015). The unexpected power of ANNs to discover unintuitive structure in the world suggests that our attempts to intuitively quantify the statistical structure in the world may fall short. How confident are we that multimodal inputs are in fact not so rich?

#### Shallow self-supervision and external-supervision are sufficient for learning

Supervision may be guided by external forces, such as social others. Even in examples of strict external supervision in machine learning, the “correct” labels are typically provided by human annotators (i.e., BNNs). In the absence of external supervision, the brain (and ANNs) can rely on self-supervised objective functions, such as prediction across space (e.g., across image patches; Doersch et al., 2015; Pathak et al., 2016), time (e.g., across video frames; Wang and Gupta, 2015; Lotter et al., 2016), or relative to self-motion or action (Agrawal et al.,. 2015; Pathak et al., 2017). In fact, in the context of prediction, the body (including adjacent computing elements in the brain) and the world itself provide abundant feedback (see Box 3). This resonates with the notion of “predictive coding” in neuroscience, which has gained momentum over the past two decades (Rao and Ballard, 1999) and is a central pillar of recent, optimization-oriented theories of brain function (Friston, 2010; Clark, 2013; Heeger, 2017).

##### Box 3. Embodiment and objective functions

Objective functions guide direct-fit optimization to generate mappings from input to output. The space of possible objective functions is large, but only a subset of objective functions will yield meaningful actions and adaptive behaviors. Currently, many ANNs are disembodied and cannot actively sample or modify their world. For example, seminal externally-supervised image classification networks (e.g., Krizhevsky et al., 2012) learn to map images to labels provided by human annotators. The affordances that emerge when learning to classify images according to 1,000 labels are very simplistic relative to the affordances of complex organisms interacting with objects in the real world. Furthermore, the brain does not have strictly-defined training and test regimes as in machine learning. While certain periods of development may be particularly critical for learning, the brain is constantly readjusting synaptic weights over the lifetime. Although we do not discuss them in depth here, end-to-end reinforcement learning models (e.g., Mnih et al., 2015) provide an appealing alternative to simplistic external supervision. In fact, the brain may adaptively shift learning strategies (e.g., from externally-supervised to self-supervised) over time.

### Direct-fit models and the school of ecological psychology

James Gibson led the school of ecological psychology, which provided an alternative account to visual perception, called direct perception, which was rejected and ultimately forgotten by many cognitive scientists. According to Gibson (1979), the brain does not aim to reconstruct the world from noisy retinal images, but rather directly detects the relevant information needed for action from a rich array of input. The school of ecological psychology did tremendous work in showing how rich the visual input is and how actions guide the selection of relevant information from the environment. However, the ecological psychology school’s critique of the traditional, strongly representational, computational approach, evoked resentment and skepticism in the field, which took the position that, without workable computational models, the argument in favor of direct perception seemed vague and unscientific (Ullman, 1980; cf. Pezzulo and Cisek, 2016). Interestingly, in a strange twist of history, advances in ANNs and the idea of direct fit provide the missing computational framework needed for the ecological school of thought. Direct fit, as an algorithmic procedure to minimize an objective function, allows neural networks to learn the transformation between external input to meaningful actions, without the need to explicitly represent underlying rules and principles in a human-interpretable way.

A major task taken up by the school of ecological psychology was to characterize each animal’s objective functions, conceptualized as affordances, based on the information the animal needs behave adaptively and survive in the world (Gibson, 1979, Michaels and Carello, 1981). For cats, a chair may afford an intermediate surface for jumping onto the kitchen counter, while for humans it may afford a surface on which to sit while eating. Like in evolution, there is no one correct way to fit the world, and different direct-fit networks, guided by different objective functions, can be used in the same ecological niche to improve fit to different aspects of the environment. Furthermore, as argued by the school of ecological psychology, information is defined as the affordances, which emerge in interaction between the organism and its ecological niche. As opposed to strongly representational approaches common in computational neuroscience, the direct-fit approach learns arbitrary functions for facilitating behavior, and is capable of mapping sensory input to motor actions without ever explicitly reconstructing the world or learning explicit rules about the latent structure of the outside world. Marr (1982), for example, speaks favorably of Gibson’s theory of vision, but, unsatisfied with the theory’s vague treatment of information processing, suggests instead that the goal of vision is to recover a geometrical representation of the world. In contrast to the representational stance, the direct-fit framework is aligned with Gibson’s treatment of the goal of vision, to recover information in the world that affords the organism its adaptive behaviors.

Gibson believed that animals are entangled with their environment in a closed perception–action feedback loop: they perceive to act and act to perceive. Furthermore, actions and affordances are shaped and constrained by the structure of the environment as well as the organism’s physiology. Similarly, from the direct-fit perspective, neural networks implicitly learn the structure of the environment as a means to an end, but this learning is ultimately driven by internal objectives aligning perception to action with an eye toward adaptive fitness (see Box 3).

### Nature versus nurture

The links between evolution and neural networks provide a fresh perspective on the nature-versus-nurture debate. So far, we have discussed how biological (and artificial) neural networks learn the structure of the world directly from examples using direct-fit optimization procedures. The ability to learn particular functions, however, is highly constrained by (1) the structure of the body, peripheral nervous system, and the properties of the sensory receptors; (2) the architecture of neural circuits; (3) the balance between pre-wired networks and open-ended plasticity. Therefore, no BNN can be considered a tabula rasa, as all three factors differ across species and are mindlessly tuned over time by evolution (Zador, 2019).

#### Bodily structure

The ability of an organism to pursue an objective function is constrained by the structure of the body. Each organism has a particular morphology (e.g., skeletal system, motor system, and sensory system) that constrains its affordances and the way in which it adapts to its ecological niche. Because evolution proceeds incrementally, the current morphology of an organism constrains the adaptations that may occur in subsequent generations (a form of regularization). Furthermore, the properties of the sensory organs constrain the type of information an organism can capitalize on. For example, bats have unique skeletal and echolocation systems, which enables their neural networks to learn how to navigate and hunt aerially in the dark. The design of the network’s peripheral structures is optimized through evolution, and though only minimally modifiable, is the backbone that shapes learning.

#### Neural circuit architecture

The ability to learn is shaped by neural circuit architecture. In contrast to Marr’s (1982) distinction between hardware and software, circuit architecture in BNNs and ANNs is tightly coupled to computation. There are many different architectures, each optimized for learning specific ad hoc tasks. For example: adding convolutional filters allows the networks to learn patterns across space (Krizhevsky et al., 2012); adding recurrent connections allows the networks to detect patterns across time (Graves, 2013); adding short- and long-term controllers allows the network to adjust the timescale over which it accumulates information (El Hihi et al., 1996; Hermans et al., 2013); adding attentional mechanisms allow the network to enhance relevant information (Luong et al., 2015; Xu et al., 2015); and adding context-based memory storage, as in differentiable neural computers, allows the network to both store episodic contexts and generalize across examples (Graves et al., 2016). Introducing novel architectural motifs is likely to improve the performance of ANNs. In BNNs, the architecture of the neural circuitry is optimized by evolution, and ranges from largely diffuse nerve nets in jellyfish, to series of ganglia in insects, to the complex subcortical and cortical structures of mammals (Satterlie, 2011; Striedter, 2005). The detailed comparative mapping of biological neural circuit architectures, learning rules, and objective functions is an active field of research, and we have much to learn from evolution’s solutions across neural systems and across organisms (Nieuwenhuys et al., 2014; Liebeskind et al., 2016).

#### Evolutionary hardwiring

In addition to optimizing the architecture of the peripheral and central components of neural networks, evolution can pre-train and optimize the synaptic weights of the networks. The retina, for example, is a specialized neural circuit optimized by evolution to convert light into neural signals and performs fairly sophisticated preprocessing on the incoming images (Carandini and Heeger, 2012). The architecture of retinal circuits is fixed (Briggman et al., 2011), and since they do not receive top-down modulation signals from cortex, the degree of neural plasticity is relatively low compared to the cortex. Similarly, many of the neuronal circuits in insect and mammalian brains are pre-wired, and ready to operate from birth (Gaier et al., 2019). Unlike other species, much of human learning takes place after birth, although some pre-trained optimization no doubt facilitates learning (Zador, 2019). Interestingly, related optimization processes like overproduction and selection may also guide plasticity in development (Changeux and Danchin, 1976; Edelman, 1993).

The parallels between evolution and learning redefine the debate on nature versus nurture. A prominent view in developmental psychology (e.g., Spelke et al., 1992; Spelke, 2007, Marcus 2018b) argues that learning relies on innate knowledge about the structure of the world (e.g., knowledge about grammar, object permanence, numerosity, etc.). In contrast, the direct-fit perspective argues that there is little need for domain-specific templates or innate, explicit knowledge of these underlying rules for the brain to function in the world (e.g., Arcaro et al., 2017). It would be inefficient to hardwire these faculties if they can be extracted from the world during development. Our affordances are constrained by our bodies and brains, and there is an intimate relationship between how our bodies and neural networks are wired and what we can learn. Framing both evolution and learning in terms of highly related optimization processes operating over different timescales mitigates the polemical character of this debate.

### At which level does psychology emerge?

We generally assume that human cognitive capacity extends beyond the “mindless” competence embedded in direct-fit models. While direct-fit models can interpolate, they lack any explicit understanding of the underlying rules and processes, which shape the world. In contrast, human cognition, at its best, provides us with tools to understand the world’s underlying structure and seek global rules; the type of understanding needed to extrapolate to qualitatively novel situations. Our minds can recombine words into new sentences, aggregate memories, and invent fictional stories. Furthermore, our minds develop mathematical and logical systems, and mechanical tools to harness knowledge and expand our capacity to understand and act in the world, capacities which seem out of reach for the direct-fit over-parameterized models.

We think that cognitive and computational neuroscience has erred in imposing extrapolation criteria and ideal-fit models wholesale on the brain. This way of thinking leverages some of the most marvelous capacities of the human mind (sometimes referred to as “System 2”; Evans et al., 1984) to explain how the brain effortlessly performs many of its cognitive tasks (referred to as “System 1”). While the human mind inspires us to touch the stars, it is grounded in the mindless billions of direct-fit parameters of System 1. Therefore, direct-fit interpolation is not the end goal, but the starting point for understanding the architecture of higher-order cognition. There is no other substrate from which System 2 could arise. Many of the processes in System 1 are shared with other animals (as in perceptual systems), and some are unique to humans (as in grammar learning), but all are executed in an automatic, fast, and often unconscious way. The brute-force direct-fit interpolation that guides learning in these systems, similar to evolution, can go further than we previously thought in explaining many cognitive functions in humans (e.g., learning syntax in natural text without imposing rule-based reasoning; see Box 2).

We still do not know the extent to which the human cognitive capacities attributed to System 2 in fact go beyond the quick and automatic procedures of System 1. Everyday, new ANN architectures are developed using direct-fit procedures to learn and perform more complex cognitive functions, such as driving, translating languages, learning calculus, or making a restaurant reservation—functions which were historically assumed to be under the jurisdiction of System 2. At the same time, these artificial networks, as opposed to humans, fail miserably in situations that require generalization and extrapolation across contexts (Lake et al., 2017). Instead of imposing efficiency, simplicity, and interpretability wholesale across neural system, psychologists should ask how our uniquely human cognitive capacities can extract explicit and compact knowledge about the outside world from the billions of direct-fit model weights. While the ability to recognize faces, speak, read, and drive may be grounded in a mindless fit to nature, our ability to abstract and verbalize information from these embeddings allows us to develop social structures, discover laws of nature, and reshape the world.

How high-level cognitive functions emerge from brute-force over-parameterized biological neural networks is likely to be a central question for future cognitive studies. Such an understanding may be necessary for developing the next generation of sentient ANNs, capable of not only sensing and acting, but also understanding and communicating the structure of the world on our terms.

## Conclusion

Historically, we have evaluated our scientific models according to low-dimensional, psychologically interpretable criteria and have thus underestimated the power of mindless over-parameterized optimization to solve complex problems in real-life contexts. We have selectively searched for explicit, low-dimensional knowledge embedded in the neural code. The expressivity of ANNs as universal approximators should be troubling to experimental neuroscientists. We typically use controlled, low-dimensional stimuli and tasks to probe brain–behavior relationships, seeking elegant, human-interpretable design principles. The analogy with evolution (and the historical argument for intelligent design) is incisive here: although intuitive design principles may emerge from neural data under experimental manipulation, these factors are incidental properties of a flexible, direct-fit learning system for modeling the natural world, and the “design” is that imposed by the experimenter.

If we evaluated ecosystems produced by evolution in terms of ideal design principles, we would find them inefficient and inscrutable. If we evaluate biological (and artificial) neural networks by the complexity of their fitted parameters, we will similarly view them as inelegant and uninterpretable. But interpretability is not strictly synonymous with elegance or simplicity. Evolutionary theory has taught us the power of mindless, iterative processes guided by natural selection to construct organisms that can navigate the world. And in fact, until fairly recently, evolution was the only such mindless process known to create self-organizing, well-adapted models of the world (Langton, 1995; Bedau, 2003). Humans have begun to create models, and simulated organisms in some cases, that, while still quite limited, can perform particular behaviors surprisingly well. It should come as no surprise that the processes required to create such models parallel evolutionary processes. The importance of evolutionary theory was in reorienting us to a previously unappreciated kind of explanation and understanding in biology.

Artificial neural networks are beginning to reveal the power of mindless overparameterized optimization guided by objective functions over a densely-sampled real-world parameter space. Despite their relative simplicity, this achievement demands we reorient our criteria for understanding biological neural networks and may require us to reevaluate the foundational assumptions of our experimental method. Using contrived experimental manipulations in hopes of recovering simple, human-interpretable representations from direct-fit neural networks—both biological and artificial—may never yield the kind of understanding we seek. The direct-fit perspective emphasizes the tight link between the structure of the world and the structure of the brain. We see a certain optimism in this view, as it provides a fresh window onto the neural code. Evolutionary theory provides a relatively simple framework for understanding an incredible diversity of phenomena; to claim that evolutionary theory is not parsimonious would be misleading. Similarly, the neural machinery that guides behavior may abide by simpler principles than our vast taxonomy of piecemeal neural representations and cognitive processes would suggest. We hope that the implications of this perspective will shine a light on the inadequacies of the reductionist approach and push the field toward more ecological, holistic approaches for studying the links between organism and environment.

While we acknowledge that ANNs are indeed highly simplified models of BNNs, we argue that there are some critical similarities: they belong to the same family of over-parameterized, direct-fit models that rely on dense sampling for learning task-relevant structures in data. In many domains, ANNs are currently the only models that attain human-like behavioral performance and they can provide unexpected insights into both the power and limitations of the direct-fit approach.

Like BNNs, ANNs are based on a collection of connected nodes called artificial neurons or units that loosely resemble the neurons in a biological nervous system. Each connection, like the synapses in BNN, links one artificial neuron to another, and the strength of these connections can be adjusted by learning. Like their biological counterparts, an artificial neuron receives signals from many neurons, integrates their input, and sends a signal to artificial neurons connected to it. The output of each artificial neuron is typically some nonlinear function of its inputs. Similarly, biological neurons typically only transmit a signal if the aggregated input signals reach a threshold. The connections between artificial neurons are assigned weights that are adjusted as learning proceeds (e.g., using the backpropagation algorithm; Rumelhart et al., 1986) based on supervised feedback or reward signals. The weight increases or decreases the strength of a connection. Similarly to BNNs, ANNs are sometimes organized into layers, and the network as a whole is optimized to map the input to the desired output according to the objective function. For additional details on the parallels between artificial and biological neural networks, we point the reader to recent reviews (Kriegeskorte, 2015; Kumaran et al., 2016; Yamins and DiCarlo, 2016; Hassabis et al., 2017; Botvinick et al., 2019; Cichy and Kaiser, 2019; Richardson et al., 2019; Whittington and Bogacz, 2019).

To make the notion of direct-fit interpolation concrete, we briefly describe two different modern ANNs: (1) a deep convolutional neural network trained to recognize faces from images using an externally-supervised objective function (Schroff et al., 2015); and (2) a transformer network that learns a language model using a self-supervised objective function (Radford et al., 2019). In both cases, rather than using engineered features, the models learn an embedding space by optimizing an objective function on densely sampled training data. Note that in both the externally-supervised case (FaceNet) and self-supervised case (GPT-2), the objective functions are ultimately governed by human behavior.

### Face model (FaceNet)

This face recognition model (Schroff et al., 2015) assumes that all facial identities in the world are embedded in a multidimensional Euclidean space (a property of the external world). While the precise number of dimensions is unknown, empirically we need the embedding space to be of sufficiently high dimension to capture all variations across individual identities. The model is supplied face images (cropped to isolate the face and represented as 220 × 220-pixel images with 3 color channels) and learns a mapping from the 145,200-dimensional pixel space to a compact 128-dimensional identity space. One of the best-performing variants of the model is a deep convolutional neural network with 22 layers (140 million parameters in total) trained using stochastic gradient descent with backpropagation. End-to-end learning is guided by an objective function (triplet loss) minimizing the distance between faces belonging to the same identity and enforcing a margin between different identities in the embedding space. This objective function effectively compresses all face images belonging to the same person into a common location in the 128-dimensional embedding space while discounting uninformative dimensions in image space and the input layers.

According to the direct-fit framework, the generalization of this model (i.e., its capacity to correctly classify face images of both familiar and novel identities) is bounded by the density and diversity of the training set (i.e., the interpolation zone in Figure 1E–F and Figure 4). If the training set spans the space of facial variability in the real world (including identity, expression, viewpoint, lighting, occlusion, etc.) with sufficiently dense examples, the model can learn an embedding space where effectively any face can be interpolated to the correct identity cluster. This superhuman, nearly perfect generalization is obtained when the network is trained on an exceptionally dense training set of 200 million face images with diverse set of 8 million identities. Importantly, the model generalizes to a test set of one million face images for novel identities not included in the training set and achieves 95–99% accuracy on common benchmark datasets. The exact same network, however, will exhibit the “other-race effect” (Malpass and Kravitz, 1969; O’Toole et al., 2018) if we will restrict the training set to, e.g., Western faces, while systematically excluding East Asian face images from the training set—thereby inducing a bias by contracting the interpolation zone. Along these lines, humans are not face experts, but rather experts in recognizing the roughly 5,000 faces they are familiar with (Jenkins et al., 2018; Young and Burton, 2018). We predict that if we selectively trained the exact same network on a cluster of 5,000 identities and a few million examples (more realistic input for the human brain), the model will learn a sparse, restricted region of identity space and will display more human-like performance. Training the same network on a restricted set of, e.g., 20 identities in a laboratory settings will result in a constrained “overfit” model capable of identifying new images sampled from within the narrow scope of this training set.

**Figure 4.**
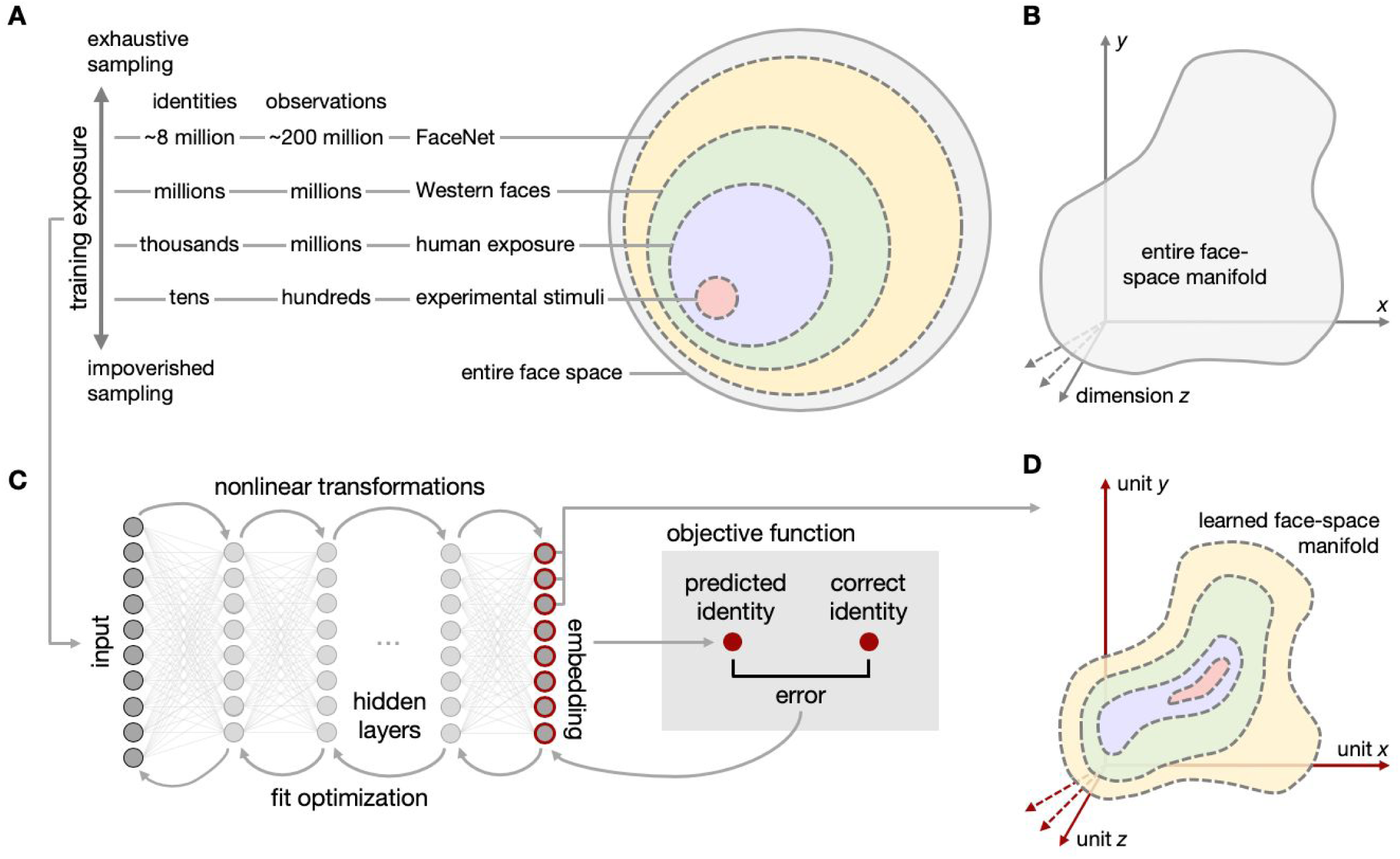
The density and diversity of training examples determines the interpolation zone and allows ANNs to approximate the regions of the face-space manifold to which they are exposed. (***A***) The scope of exposure may range from: controlled experimental stimuli (e.g., Guntupalli et al., 2016); to typical human exposure (Jenkins et al., 2018); to a biased sample of only Western faces (O’Toole et al., 2018); to the vast training sample supplied to FaceNet (Schroff et al., 2015). All of these are subsets of the entire face space. Note that the numbers of identities and observations indicated in panel A are crude approximations for illustrative purposes. (***B***) All facial variation in the world can be represented geometrically as locations on a manifold in an abstract, high-dimensional “face space” constrained by the physical properties of human physiognomy. (***C***) A simple schematic depiction of an ANN, which maps input images (e.g., pixel values for face images) through many hidden layers into a lower-dimensional embedding space. The network’s objective function quantifies the mismatch between the model’s output and the desired output (e.g., predicting the correct identity). Error signals are then propagated back through the network to adjust connection weights, incrementally optimizing the network to better perform the task specified by the objective function within the boundaries of the training data (interpolation zone). Note that modern ANNs have drastically more complex architectures than depicted in the schematic (e.g., convolutional layers). (***D***) Training an ANN such as FaceNet (Schroff et al., 2015) on a vast number of diverse face images yields an interpolation zone encompassing nearly all facial variation in the world (yellow; superhuman performance). However, training the exact same model on only Western faces will yield a constrained interpolation zone (green), and the model will generalize poorly to faces outside this interpolation zone (the “other-race effect”). When trained on a sparser sample representative of typical human exposure, the network will yield human-like performance (purple). Finally, if trained on impoverished data, the model will nonetheless interpolate well within the scope of this limited training set, but will fail to generalize beyond. The interpolation zone is a result of the density and diversity of the training sample.

A few lessons become clear from this example: generalization is bounded by the interpolation zone, which is determined by properties of the training set (i.e., density and diversity). The difficulty of the learning task is constrained by the complexity of the task-relevant manifold on which the data reside as approximated by the multidimensional embedding space (e.g., a continuous, smooth, low-dimensional manifold may facilitate learning). Note that these are properties of the external world (as expressed in the training set) and not strictly properties of the network. Focusing exclusively on interpreting the properties of the 128-dimensional embedding layer can be misleading for several reasons. First, the embedding layer is the tip of the iceberg: the embedding space is the result of an over-parameterized direct-fit learning process, and the behavioral performance of the model is the joint product of the architecture, objective function, learning rule, training set, and so on; we cannot ignore the training sample or the computational motifs that yield the embedding space if we hope to understand how the neural network works (for related arguments, see Jonas and Kording, 2017; Lillicrap and Kording, 2019; Richards et al., 2019). Second, in the context of direct-fit learning with exhaustive sampling, the structure of the embedding space generally reflects the task-relevant structure of the external world. We should exercise caution in interpreting particular structural properties of the embedding space as “intrinsic” properties of the network. Finally, given the multidimensionality of real-life input (e.g., 145,200-dimensional pixel inputs for FaceNet) and the multidimensionality of the face-space manifold in the world (e.g., 100+ dimensions), the program of running highly-controlled experiments in an attempt to find low-dimensional, psychologically-interpretable neural response features may lead us astray.

### Language model (GPT-2)

This language model (Radford et al., 2019) assumes that there are sufficient regularities in the way people use language in specific contexts to learn a variety of complex linguistic tasks (Wittgenstein, 1953). Again, we emphasize that the quality of the model will be constrained by the density and diversity of examples provided during training. Specifically, the model uses an attention-based “transformer” architecture (Vaswani et al., 2017) with 48 layers and over 1.5 billion parameters to perform sequence transduction. In simple terms, the transformer can be thought of as a coupled encoder and decoder where the input to the decoder is shifted to the subsequent element (i.e., the next word or byte). Critically, both the encoder and decoder components are able to selectively attend to elements at nearby positions in the sequence, effectively incorporating contextual information. The model is trained on over 8 million documents for a total of 40 gigabytes of text. Despite the self-supervised sequence-to-sequence learning objective, the model excelled at a variety of linguistic tasks, such as predicting the final word of lengthy sentences, question answering, summarization, and translation, approaching human performance in some cases. Contextual prediction is a cognitively appealing self-supervised objective function, as it is readily available to all learners at all stages of development. Furthermore, the self-supervised objective function is still shaped by external human behaviors in real-life contexts, which provide a structured linguistic input to the learner, exposing the entwined relationships between self-supervision and external-supervision. As opposed to humans, however, this model only learns to predict based on a relatively narrow behavioral context (the preceding words), and is deprived of actions, corroborating visual cues (cf. Vinyals et al., 2015), and social cues available to humans (see Box 3). The capacity to learn temporal dependencies over many words still does not compare to our ability to accumulate and integrate broadly-distributed multimodal information over hours, days, and years. Despite the limitation of the training set and objective function, surprisingly, models of this kind (e.g., Devlin et al., 2018) may also implicitly learn some compositional properties of language, such as syntax, from the structure of the input (Linzen et al., 2016; Belinkov et al., 2017; Baroni, 2019; Hewitt and Manning, 2019).

Objective functions in BNNs must also satisfy certain constraints imposed by the body to behave adaptively when interacting with the world. Examples of objective functions guided by action include: learning to balance the body while walking across the room, learning to coordinate hands and eyes to touch objects, and learning to coordinate hand and finger movements to bring food to the mouth. In all these cases, it is clear whether the brain succeeded or failed at each trial, and it is clear how minimizing cost functions can provide the necessary feedback to guide the fit without appealing to an explicit, rule-based understanding of the physical forces at work. Furthermore, analogous to the gradual innovation along the evolutionary tree, in which a new function is scaffolded by prior advances, learning one objective function, such as standing, paves the way for learning a new objective function, such as walking, which can further enable running, jumping, or dancing.

Another source of guidance for learning is the actions of other agents within the social network. Examples of objective functions guided by other brains include learning to recognize individual faces, learning to name objects, learning to produce grammatical sentences, and learning to read. In all of these examples, the solution is provided by social others (Wittgenstein, 1953, Hasson et al., 2012). Because social exchange provides a basis for external supervision, the brain can rapidly learn complex knowledge collectively accumulated over generations. Therefore, adding to current ANNs a body that is capable of actively sampling and interacting with the world (e.g., Levine et al., 2018), as well as adding means to directly interact with other networks (e.g., Goodfellow et al., 2014; Jaderberg et al., 2019), may increase the network’s capacity to learn and reduce the gaps between BNNs and ANNs (Marblestone et al., 2016; Baker et al., 2019; Leibo et al., 2019).

## Acknowledgements

We thank Hezy Yeshurun, Galia Avidan, Eran Malach, Bruno Galantucci, Asif A. Ghazanfar, Rita Goldstein, Liat Hasenfratz, Hanna Hillman, Meir Meshulam, Christopher J. Honey, Rafael Malach, Qihong Lu, and Kenneth A. Norman for extensive discussion and helpful comments on the manuscript. This work was supported by the National Institutes of Health under award numbers DP1HD091948 (U.H., A.G.) and R01MH112566-01 (S.A.N.).

## Footnotes

1 “… the sciences do not try to explain, they hardly even try to interpret, they mainly make models. By a model is meant a mathematical construct which, with the addition of certain verbal interpretations, describes observed phenomena. The justification of such a mathematical construct is solely and precisely that it is expected to work—that is, correctly to describe phenomena from a reasonably wide area.”

2 We use the terms “interpolation”, “extrapolation”, and “robust” more broadly than their strict mathematical usage.

## References

Adami, C. (2012). The use of information theory in evolutionary biology. Ann. N. Y. Acad. Sci. 1256, 49–65.

Agrawal, P., Carreira, J., and Malik, J. (2015). Learning to see by moving. In Proc. IEEE Int. Conf. Comput. Vis., pp. 37–45.

Anderson, M., and Chemero, A. (2016). The brain evolved to guide action. In The Wiley Handbook of Evolutionary Neuroscience, S. V. Shepherd, ed. (Chichester, England: John Wiley and Sons), pp. 1–20.

Arcaro, M. J., Schade, P. F., Vincent, J. L., Ponce, C. R., and Livingstone, M. S. (2017). Seeing faces is necessary for face-domain formation. Nat. Neurosci. 20, 1404–1412.

Ashby, W. R. (1956). An Introduction to Cybernetics (London, England: Chapman and Hall).

Azevedo, F. A., Carvalho, L. R., Grinberg, L. T., Farfel, J. M., Ferretti, R. E., Leite, R. E., Filho, W. J., Lent, R., and Herculano-Houzel, S. (2009). Equal numbers of neuronal and nonneuronal cells make the human brain an isometrically scaled-up primate brain. J. Comp. Neurol. 513, 532–541.

Bäck, T. (1996). Evolutionary Algorithms in Theory and Practice: Evolution Strategies, Evolutionary Programming, Genetic Algorithms (Oxford, England: Oxford University Press).

Baker, B., Kanitscheider, I., Markov, T., Wu, Y., Powell, G., McGrew, B., and Mordatch, I. (2019). Emergent tool use from multi-agent autocurricula. arXiv, 1909.07528.

Bansal, Y., Advani, M., Cox, D. D., and Saxe, A. M. (2018). Minnorm training: an algorithm for training over-parameterized deep neural networks. arXiv, 1806.00730.

Baroni, M. (2019). Linguistic generalization and compositionality in modern artificial neural networks. arXiv, 1904.00157.

Bedau, M. A. (2003). Artificial life: organization, adaptation and complexity from the bottom up. Trends Cogn. Sci. 7, 505–512.

Belinkov, Y., Durrani, N., Dalvi, F., Sajjad, H., and Glass, J. (2017). What do neural machine translation models learn about morphology? arXiv, 1704.03471.

Botvinick, M., Ritter, S., Wang, J. X., Kurth-Nelson, Z., Blundell, C., and Hassabis, D. (2019). Reinforcement learning, fast and slow. Trends Cogn. Sci. 23, 408–422.

Breiman, L. (2001). Statistical modeling: the two cultures. Stat. Sci. 16, 199–231.

Briggman, K. L., Helmstaedter, M., and Denk, W. (2011). Wiring specificity in the direction-selectivity circuit of the retina. Nature 471, 183–188.

Brunswik, E. (1947). Perception and the Representative Design of Psychological Experiments (Berkeley, CA: University of California Press).

Carandini, M., and Heeger, D. J. (2012). Normalization as a canonical neural computation. Nat. Rev. Neurosci. 13, 51–62.

Changeux, J. P., and Danchin, A. (1976). Selective stabilisation of developing synapses as a mechanism for the specification of neuronal networks. Nature 264, 705–712.

Chomsky, N. (1965). Aspects of the Theory of Syntax (Cambridge, MA: MIT Press).

Chomsky, N. (1980). Rules and Representations (New York, NY: Columbia University Press).

Christiansen, M. H., and Chater, N. (2008). Language as shaped by the brain. Behav. Brain Sci. 31, 489–509.

Cichy, R. M., and Kaiser, D. (2019). Deep neural networks as scientific models. Trends Cogn. Sci. 23, 305–317.

Clark, A. (2013). Whatever next? Predictive brains, situated agents, and the future of cognitive science. Behav. Brain Sci. 36, 181–204.

Conant, R. C., and Ross Ashby, W. (1970). Every good regulator of a system must be a model of that system. Int. J. Syst. Sci. 1, 89–97.

Cybenko, G. (1989). Approximation by superpositions of a sigmoidal function. Math. Control Signals Syst. 2, 303–314.

Darwin, C. (1859). On the Origin of Species (London, England: John Murray).

Dawkins, R. (1986). The Blind Watchmaker: Why the Evidence of Evolution Reveals a Universe Without Design (New York, NY: Norton).

Dennett, D. C. (1995). Darwin’s Dangerous Idea: Evolution and the Meanings of Life (New York, NY: Simon and Schuster).

Dennett, D. C. (2017). From Bacteria to Bach and Back: The Evolution of Minds (New York, NY: Norton).

Devlin, J., Chang, M. W., Lee, K., and Toutanova, K. (2018). BERT: pre-training of deep bidirectional transformers for language understanding. arXiv, 1810.04805.

Doersch, C., Gupta, A., and Efros, A. A. (2015). Unsupervised visual representation learning by context prediction. In Proc. IEEE Int. Conf. Comput. Vis., pp. 1422–1430.

Edelman, G. M. (1993). Neural Darwinism: selection and reentrant signaling in higher brain function. Neuron 10, 115–125.

Eiben, A. E., and Smith, J. (2015). From evolutionary computation to the evolution of things. Nature 521, 476–482.

El Hihi, S., and Bengio, Y. (1996). Hierarchical recurrent neural networks for long-term dependencies. In Adv. Neural Inf. Process. Syst., pp. 493–499.

Eliasmith, C., Stewart, T. C., Choo, X., Bekolay, T., DeWolf, T., Tang, Y., and Rasmussen, D. (2012). A large-scale model of the functioning brain. Science 338, 1202–1205.

Evans, J. S. B. (1984). Heuristic and analytic processes in reasoning. Bri. J. Psychol. 75, 451–468.

Feldman, V. (2019). Does learning require memorization? A short tale about a long tail. arXiv, 1906.05271.

Felsen, G., and Dan, Y. (2005). A natural approach to studying vision. Nat. Neurosci. 8, 1643–1646.

Felleman, D. J., and Van Essen, D. C. (1991). Distributed hierarchical processing in the primate cerebral cortex. Cereb. Cortex 1, 1–47.

Fisher, R. A. (1930). The Genetical Theory of Natural Selection (Oxford, England: Clarendon Press).

Fisher, R. A. (1935). The Design of Experiments (Edinburgh, England: Oliver and Boyd).

Fodor, J. A. (1983). Modularity of Mind: An Essay on Faculty Psychology (Cambridge, MA: MIT Press).

Friston, K. (2010). The free-energy principle: a unified brain theory? Nat. Rev. Neurosci. 11, 127–138.

Funahashi, K. I. (1989). On the approximate realization of continuous mappings by neural networks. Neural Netw. 2, 183–192.

Gaier, A., and Ha, D. (2019). Weight agnostic neural networks. arXiv, 1906.04358.

Gao, P., Trautmann, E., Byron, M. Y., Santhanam, G., Ryu, S., Shenoy, K., and Ganguli, S. (2017). A theory of multineuronal dimensionality, dynamics and measurement. bioRxiv, 214262.

Gibson, J.J. (1979). The Ecological Approach to Visual Perception (Boston, MA: Houghton Mifflin).

Godfrey-Smith, P. (2007). Information in biology. In The Cambridge Companion to the Philosophy of Biology, D. Hull, and M. Ruse, eds. (Cambridge, England: Cambridge University Press), pp. 103–119.

Gomez-Marin, A., and Ghazanfar, A. A. (2019). The life of behavior. Neuron 104, 25–36.

Goodfellow, I., Pouget-Abadie, J., Mirza, M., Xu, B., Warde-Farley, D., Ozair, S., Courville, A., and Bengio, Y. (2014). Generative adversarial nets. In Adv. Neural Inf. Process. Syst., pp. 2672–2680.

Gould, S. J. (1982). Darwinism and the expansion of evolutionary theory. Science 216, 380–387.

Gould, S. J., and Lewontin, R. C. (1979). The spandrels of San Marco and the Panglossian paradigm: a critique of the adaptationist programme. Proc. R. Soc. Lond. B. Biol. Sci. 205, 581–598.

Graves, A., Wayne, G., Reynolds, M., Harley, T., Danihelka, I., Grabska-Barwińska, A., Colmenarejo, S. G., Zwols, Y., Ostrovski, G., Cain, A. et al. (2016). Hybrid computing using a neural network with dynamic external memory. Nature 538, 471–476.

Graves, A., Mohamed, A. R., and Hinton, G. (2013). Speech recognition with deep recurrent neural networks. In Proc. IEEE Int. Conf. Acoust. Speech Signal Process., pp. 6645–6649.

Guntupalli, J. S., Wheeler, K. G., and Gobbini, M. I. (2016). Disentangling the representation of identity from head view along the human face processing pathway. Cereb. Cortex 27, 46–53.

Hamilton, L. S., and Huth, A. G. (2018). The revolution will not be controlled: natural stimuli in speech neuroscience. Lang. Cogn. Neurosci.

Hassabis, D., Kumaran, D., Summerfield, C., and Botvinick, M. (2017). Neuroscience-inspired artificial intelligence. Neuron 95, 245–258.

Hasson, U., Ghazanfar, A. A., Galantucci, B., Garrod, S., and Keysers, C. (2012). Brain-to-brain coupling: a mechanism for creating and sharing a social world. Trends Cogn. Sci. 16, 114–121.

Hasson, U., and Honey, C. J. (2012). Future trends in neuroimaging: neural processes as expressed within real-life contexts. NeuroImage 62, 1272–1278.

Heeger, D. J. (2017). Theory of cortical function. Proc. Natl. Acad. Sci. U.S.A. 114, 1773–1782.

Hermans, M., and Schrauwen, B. (2013). Training and analysing deep recurrent neural networks. In Adv. Neural Inf. Process. Syst., pp. 190–198.

Hewitt, J., and Manning, C. D. (2019). A structural probe for finding syntax in word representations. In Proc. sNorth Am. Chap. Assoc. Comput. Linguist. Hum. Lang. Technol., pp. 4129–4138.

Hinton, G. E., Mcclelland, J. L., and Rumelhart, D. E. (1986). Distributed representations. In Parallel Distributed Processing: Explorations in the Microstructure of Cognition, Vol. 1: Foundations, D. E. Rumelhart, J. L. McClelland, and The PDP Research Group, eds. (Cambridge, MA: MIT Press), pp. 77–109.

Hochreiter, S., and Schmidhuber, J. (1997). Long short-term memory. Neural Comput. 9, 1735–1780.

Holland, J. H. (1992). Adaptation in Natural and Artificial Systems: An Introductory Analysis with Applications to Biology, Control and Artificial Intelligence (Ann Arbor, MI: University of Michigan Press).

Hornik, K., Stinchcombe, M., and White, H. (1989). Multilayer feedforward networks are universal approximators. Neural Netw. 2, 359–366.

Hubel, D. H., and Wiesel, T. N. (1962). Receptive fields, binocular interaction and functional architecture in the cat’s visual cortex. J. Physiol. 160, 106–154.

Jaderberg, M., Czarnecki, W. M., Dunning, I., Marris, L., Lever, G., Castañeda, A. G., Beattie, C., Rabinowitz, N. C., Morcos, A. S., Ruderman, A. et al. (2019). Human-level performance in 3D multiplayer games with population-based reinforcement learning. Science 364, 859–865.

Jenkins, R., Dowsett, A. J., and Burton, A. M. (2018). How many faces do people know? Proc. R. Soc. Lond. B. Biol. Sci. 285, 20181319.

Jolly, E., and Chang, L. J. (2019). The Flatland fallacy: moving beyond low-dimensional thinking. Top. Cogn. Sci. 11, 433–454.

Jonas, E., and Kording, K. P. (2017). Could a neuroscientist understand a microprocessor? PLoS Comp. Biol. 13, e1005268

Kandel, E. R., Schwartz, J. H., Jessell, T. M., Siegelbaum, S., Hudspeth, A. J., and Mack, S. (2012). Principles of Neural Science, 5th ed. (New York, NY: McGraw-Hill).

Krakauer, J. W., Ghazanfar, A. A., Gomez-Marin, A., MacIver, M. A., and Poeppel, D. (2017). Neuroscience needs behavior: correcting a reductionist bias. Neuron 93, 480–490.

Kriegeskorte, N. (2015). Deep neural networks: a new framework for modeling biological vision and brain information processing. Ann. Rev. Vis. Sci. 1, 417–446.

Krizhevsky, A., Sutskever, I., and Hinton, G. E. (2012). ImageNet classification with deep convolutional neural networks. In Adv. Neural Inf. Process. Syst., pp. 1097–1105.

Kumaran, D., Hassabis, D., and McClelland, J. L. (2016). What learning systems do intelligent agents need? Complementary learning systems theory updated. Trends Cogn. Sci. 20, 512–534.

Lake, B. M., Ullman, T. D., Tenenbaum, J. B., and Gershman, S. J. (2017). Building machines that learn and think like people. Behav. Brain Sci. 40, e253.

Langton, C. G. (1995). Artificial Life: An Overview (Cambridge, MA: MIT Press).

LeCun, Y., Bengio, Y., and Hinton, G. (2015). Deep learning. Nature 521, 436–444.

Leibo, J. Z., Hughes, E., Lanctot, M., and Graepel, T. (2019). Autocurricula and the emergence of innovation from social interaction: a manifesto for multi-agent intelligence research. arXiv, 1903.00742.

Levine, S., Pastor, P., Krizhevsky, A., Ibarz, J., and Quillen, D. (2018). Learning hand-eye coordination for robotic grasping with deep learning and large-scale data collection. Int. J. Rob. Res. 37, 421–436.

Levy, I., Hasson, U., and Malach, R. (2004). One picture is worth at least a million neurons. Curr. Biol. 14, 996–1001.

Lewontin, R. C. (1970). The units of selection. Ann. Rev. Ecol. Syst. 1, 1–18.

Liebeskind, B. J., Hillis, D. M., Zakon, H. H., and Hofmann, H. A. (2016). Complex homology and the evolution of nervous systems. Trends Ecol. Evol. 31, 127–135.

Lillicrap, T. P., and Kording, K. P. (2019). What does it mean to understand a neural network? arXiv, 1907.06374.

Linzen, T., Dupoux, E., and Goldberg, Y. (2016). Assessing the ability of LSTMs to learn syntax-sensitive dependencies. Trans. Assoc. Comput. Linguist. 4, 521–535.

Lotter, W., Kreiman, G., and Cox, D. (2016). Deep predictive coding networks for video prediction and unsupervised learning. arXiv, 1605.08104.

Luong, M. T., Pham, H., and Manning, C. D. (2015). Effective approaches to attention-based neural machine translation. arXiv, 1508.04025.

Malpass, R. S., and Kravitz, J. (1969). Recognition for faces of own and other race. J. Pers. Soc. Psychol. 13, 330–334.

Marblestone, A. H., Wayne, G., and Kording, K. P. (2016). Toward an integration of deep learning and neuroscience. Front. Comput. Neurosci. 10, 94.

Marcus, G. (2018a). Deep learning: a critical appraisal. arXiv, 1801.00631.

Marcus, G. (2018b). Innateness, AlphaZero, and artificial intelligence. arXiv, 1801.05667.

Marom, S., Meir, R., Braun, E., Gal, A., Kermany, E., and Eytan, D. (2009). On the precarious path of reverse neuro-engineering. Front. Comput. Neurosci. 3, 5.

Marr, D. (1982) Vision: A Computational Investigation into the Human Representation and Processing of Visual Information (San Francisco, CA: Freeman).

Maynard Smith, J. (2000). The concept of information in biology. Philos. Sci. 67, 177–194.

Mayr, E. (1992). The idea of teleology. J. Hist. Ideas 53, 117–135.

McClelland, J. L., and Rogers, T. T. (2003). The parallel distributed processing approach to semantic cognition. Nat. Rev. Neurosci. 4, 310–322.

McCloskey, M. (1991). Networks and theories: the place of connectionism in cognitive science. Psychol. Sci. 2, 387–395.

Meehl, P. E. (1990). Why summaries of research on psychological theories are often uninterpretable. Psychol. Rep. 66, 195–244.

Michaels, C. F., and Carello, C. (1981). Direct Perception (Englewood Cliffs, NJ: Prentice-Hall).

Mnih, V., Kavukcuoglu, K., Silver, D., Rusu, A. A., Veness, J., Bellemare, M. G., Graves, A., Riedmiller, M., Fidjeland, A. K., Ostrovski, G. et al. (2015). Human-level control through deep reinforcement learning. Nature 518, 529–533.

Musslick, S., Saxe, A., Özcimder, K., Dey, B., Henselman, G., and Cohen, J. D. (2017). Multitasking capability versus learning efficiency in neural network architectures. In Proc. Annu. Conf. Cogn. Sci. Soc., pp. 829–834.

Nieuwenhuys, R., Hans, J., and Nicholson, C. (2014). The Central Nervous System of Vertebrates (Berlin, Germany: Springer).

Olshausen, B. A., and Field, D. J. (2005). How close are we to understanding V1? Neural Comput. 17, 1665–1699.

O’Toole, A. J., Castillo, C. D., Parde, C. J., Hill, M. Q., and Chellappa, R. (2018). Face space representations in deep convolutional neural networks. Trends Cogn. Sci. 22, 794–809.

Pathak, D., Agrawal, P., Efros, A. A., and Darrell, T. (2017). Curiosity-driven exploration by self-supervised prediction. In Proc. IEEE Conf. Comp. Vis. Pattern Recognit. Workshops, pp. 16–17.

Pathak, D., Krahenbuhl, P., Donahue, J., Darrell, T., and Efros, A. A. (2016). Context encoders: Feature learning by inpainting. In Proc. IEEE Conf. Comp. Vis. Pattern Recognit. Workshops, pp. 2536–2544.

Pezzulo, G., and Cisek, P. (2016). Navigating the affordance landscape: feedback control as a process model of behavior and cognition. Trends Cogn. Sci. 20, 414–424.

Pinker, S. (1994). The Language Instinct: How the Mind Creates Language (New York, NY: William Morrow).

Ponce, C. R., Xiao, W., Schade, P. F., Hartmann, T. S., Kreiman, G., and Livingstone, M. S. (2019). Evolving images for visual neurons using a deep generative network reveals coding principles and neuronal preferences. Cell 177, 999–1009.

Pullum, G. K., and Scholz, B. C. (2002). Empirical assessment of stimulus poverty arguments. Linguist. Rev. 18, 9–50.

Quackenbush, J. (2001). Computational analysis of microarray data. Nat. Rev. Genet. 2, 418–427.

Radford, A., Wu, J., Child, R., Luan, D., Amodei, D., and Sutskever, I. (2019). Language models are unsupervised multitask learners.

Raffel, C., Shazeer, N., Roberts, A., Lee, K., Narang, S., Matena, M., Zhou, Y., Li, W., and Liu, P. J. (2019). Exploring the limits of transfer learning with a unified text-to-text transformer. arXiv, 1910.10683.

Raghu, M., Poole, B., Kleinberg, J., Ganguli, S., and Dickstein, J. S. (2017). On the expressive power of deep neural networks. In Proc. Int. Conf. Mach. Learn., pp. 2847–2854.

Ramon, M., and Gobbini, M. I. (2018). Familiarity matters: a review on prioritized processing of personally familiar faces. Vis. Cogn. 26, 179–195.

Ramscar, M., and Yarlett, D. (2007). Linguistic self-correction in the absence of feedback: a new approach to the logical problem of language acquisition. Cogn. Sci. 31, 927–960.

Rao, R. P. N., and Ballard, D. H. (1999). Predictive coding in the visual cortex: a functional interpretation of some extra-classical receptive-field effects. Nat. Neurosci. 2, 79–87.

Richards, B. A., Lillicrap, T. P., Beaudoin, P., Bengio, Y., Bogacz, R., Christensen, A., Clopath, C., Costa, R. P., de Berker, A., Ganguli, S. et al. (2019). A deep learning framework for neuroscience. Nat. Neurosci. 22, 1761–1770.

Roy, B. C., Frank, M. C., DeCamp, P., Miller, M., and Roy, D. (2015). Predicting the birth of a spoken word. Proc. Natl. Acad. Sci. U.S.A. 112, 12663–12668.

Rozenblit, L., and Keil, F. (2002). The misunderstood limits of folk science: an illusion of explanatory depth. Cogn. Sci. 26, 521–562.

Rumelhart, D. E., Hinton, G. E., and Williams, R. J. (1986). Learning representations by back-propagating errors. Nature 323, 533–536.

Rumelhart, D. E., McClelland, J. L., and the PDP Research Group. (1986). Parallel Distributed Processing: Explorations in the Microstructure of Cognition, Volume 1: Foundations (Cambridge, MA: MIT Press).

Satterlie, R. A. (2011). Do jellyfish have central nervous systems? J. Exp. Biol. 214, 1215–1223.

Schroff, F., Kalenichenko, D., and Philbin, J. (2015). FaceNet: a unified embedding for face recognition and clustering. In Proc. IEEE Conf. Comp. Vis. Pattern Recognit., pp. 815–823).

Srivastava, M., and Grill-Spector, K. (2018). The effect of learning strategy versus inherent architecture properties on the ability of convolutional neural networks to develop transformation invariance. arXiv, 1810.13128

Shmueli, G. (2010). To explain or to predict? Stat. Sci. 25, 289–310.

Sivak, M. (1996). The information that drivers use: is it indeed 90% visual? Perception 25, 1081–1089.

Spelke, E. S., Breinlinger, K., Macomber, J., and Jacobson, K. (1992). Origins of knowledge. Psychol. Rev. 99, 605–632.

Spelke, E. S., and Kinzler, K. D. (2007). Core knowledge. Dev. Sci. 10, 89–96.

Striedter, G. F. (2005). Principles of Brain Evolution (Sunderland, MA: Sinauer Associates).

Stringer, C., Pachitariu, M., Steinmetz, N., Carandini, M., and Harris, K. D. (2019). High-dimensional geometry of population responses in visual cortex. Nature 571, 361–365.

Taigman, Y., Yang, M., Ranzato, M. A., and Wolf, L. (2014). DeepFace: closing the gap to human-level performance in face verification. In Proc. IEEE Conf. Comp. Vis. Pattern Recognit., pp. 1701–1708.

Thompson, J. N. (2013). Relentless Evolution (Chicago, IL: University of Chicago Press).

Ullman, S. (1980). Against direct perception. Behav. Brain Sci. 3, 373–381.

Vaswani, A., Shazeer, N., Parmar, N., Uszkoreit, J., Jones, L., Gomez, A. N., Kaiser, L., and Polosukhin, I. (2017). Attention is all you need. In Adv. Neural Inf. Process. Syst., pp. 5998–6008.

Vinyals, O., Toshev, A., Bengio, S., and Erhan, D. (2015). Show and tell: a neural image caption generator. In Proc. IEEE Conf. Comp. Vis. Pattern Recognit., pp. 3156–3164.

von Neumann, J. (1955). Method in the physical sciences. In The Unity of Knowledge, L. G. Leary, ed. (Garden City, NY: Doubleday), p. 157.

Wang, X., and Gupta, A. (2015). Unsupervised learning of visual representations using videos. In Proc. IEEE Int. Conf. Comput. Vis., pp. 2794–2802.

Whittington, J. C., and Bogacz, R. (2019). Theories of error back-propagation in the brain. Trends Cogn. Sci. 23, 235–250.

Williams, G. C. (1966). Adaptation and Natural Selection: A Critique of Some Current Evolutionary Thought (Princeton, NJ: Princeton University Press).

Wittgenstein, L. (1953). Philosophical Investigations, G. E. M. Anscombe, trans. (London, England: McMillan).

Wolpert, D. H., and Macready, W. G. (1997). No free lunch theorems for optimization. IEEE Trans. Evol. Comput. 1, 67–82.

Yamins, D. L., and DiCarlo, J. J. (2016). Using goal-driven deep learning models to understand sensory cortex. Nat. Neurosci. 19, 356–365.

Yarkoni, T., and Westfall, J. (2017). Choosing prediction over explanation in psychology: lessons from machine learning. Perspect. Psychol. Sci. 12, 1100–1122.

Young, A. W., and Burton, A. M. (2018). Are we face experts? Trends Cogn. Sci. 22, 100–110.

Xu, K., Ba, J., Kiros, R., Cho, K., Courville, A., Salakhutdinov, R., Zemel, R., and Bengio, Y. (2015). Show, attend and tell: neural image caption generation with visual attention. In Proc. Int. Conf. Mach. Learn., pp. 2048–2057.

Zador, A. M. (2019). A critique of pure learning and what artificial neural networks can learn from animal brains. Nat. Commun. 10, 3770.

Zhang, C., Bengio, S., Hardt, M., Recht, B., and Vinyals, O. (2016). Understanding deep learning requires rethinking generalization. arXiv, 1611.03530.

Zyzik, E. (2009). The role of input revisited: nativist versus usage-based models. L2 J. 1, 42–61.

